# Primary bovine white blood cells support dissemination of Lumpy Skin Disease Virus while suppressing viral replication

**DOI:** 10.1101/2024.07.18.604162

**Authors:** Manoj Kumar, Ohad Frid, Asaf Sol, Alexander Rouvinski, Sharon Karniely

## Abstract

Lumpy skin disease (LSD) is a severe infectious, emerging transboundary disease of cattle, caused by a Pox family DNA virus. Lumpy skin disease virus (LSDV) infection is associated with a febrile response followed by emergence of widespread dermal nodules. In addition to the skin, LSDV resides in multiple internal organs and can be isolated from the blood of infected cattle. LSDV is suggested to be mechanically transmitted by biting arthropods. Live attenuated vaccines are commonly used to control disease and its spread. We have characterized the tropism, replication, and dissemination of a LSDV field isolate and of an attenuated vaccine strain using *in vitro* systems. To follow virus infection and dissemination in living cells, we have generated recombinant viruses expressing green fluorescent protein (GFP) under a synthetic viral promoter. Recombinant, GFP-expressing, LSDVs demonstrated similar replication kinetics to their corresponding parental LSDV strains in a bovine kidney cell line (MDBK). We further demonstrated that LSDV-GFP productively replicated in a bovine macrophage cell line and in primary bovine foreskin cells with no apparent differences between the field isolate and the vaccine strain. When bovine peripheral blood mononuclear cells (PBMCs) were infected with either LSDV recombinant strain, we observed specific viral driven GFP fluorescence as well as significant viral gene expression. However, infected PBMCs failed to support substantial viral DNA replication and release of infectious progeny. Subsequent analysis of the anti-viral response revealed that heat treated (HT) LSDV induced the expression of interferon-stimulated genes (ISGs) in PBMCs, but this response was suppressed by infectious viruses. Finally, we show that despite failed replication, LSDV infected PBMCs transmitted the virus to recipient co-cultured MDBK cells and produced infectious foci, suggesting a potential role of PBMCs in LSDV dissemination.

**Highlights:**

- Virulent and attenuated LSDV productively replicated in bovine kidney and bovine macrophage cell lines as well as in primary fibroblasts.
- Adherent white blood cells were susceptible to LSDV field and attenuated vaccine infection.
- LSDV showed active viral transcription in PBMCs yet no significant viral genome replication or production of infectious progeny.
- PBMCs infected with heat-treated LSDV but not with fully infectious viruses upregulated ISGs’ RNA.
- PBMCs transmitted and disseminated LSDV to contacting permissive cells.

## Introduction

Lumpy skin disease (LSD) is a severe, emerging transboundary disease of cattle that causes great losses to the farm industry (Tuppurainen and Oura, 2012). LSD is caused by a Pox family, enveloped DNA virus of ∼150 kb genome, encoding 156 putative open reading frames (Tulman et al., 2001). Lumpy skin disease virus (LSDV) has high genome sequence identity with the two other members of the Capripox genus, Sheeppox virus (SPPV) and Goatpox virus (GTPV) (Tulman et al., 2002). These Capripox viruses also share a high degree of antigenic resemblance and are indistinguishable from LSDV in serological assays, yet LSDV sustains high host specificity (Black et al., 1986; Hamdi et al., 2020; Tulman et al., 2001; Tuppurainen et al., 2018a). Skin nodules formation, fever, and swollen lymph nodes are characteristics of LSDV infection and are associated with reduced milk production, infertility, and abortions (Davies, 1991). LSD is endemic in many African and Middle Eastern countries and has caused several outbreaks in Europe over the past decade (Anwar et al., 2022; Beard, 2016; Thomas, 1945). In recent years, LSD is spreading in East Asian countries (Khatri et al., 2023; Ratyotha et al., 2022; Ren et al., 2023; Smaraki et al., 2024) raising great concern about its further expansion in developing nations. LSD has been enlisted as a notifiable disease by the World Organization for Animal Health (WOAH), since reoccurring outbreaks are threatening live stockbreeding and mounting economic burden on farmers (Tuppurainen and Oura, 2012; WOAH, 2023).

Transmission of LSDV is suggested to be mediated mainly by blood-feeding arthropods and their seasonal swarming can expand the outbreak’s radius (Chihota et al., 2001, 2003; Issimov et al., 2020; Suwankitwat et al., 2023; Tuppurainen et al., 2018b; Tuppurainen et al., 2013; Yeruham et al., 1995). However, there were few reports of direct transmission of the virus between host animals (Aleksandr et al., 2020). Similar to other Poxvirus diseases, immune control of LSD involves cell-mediated immunity, which requires vaccination with live attenuated viruses. This led to the use of either SPPV and GTPV heterologous viruses or an attenuated LSDV homologous virus in vaccines (Ben-Gera et al., 2015; Kitching, 2003). The LSDV attenuated (Neethling) vaccine was developed by multiple passaging of a field isolate in cultured cells and in embryonated chicken eggs (van Rooyen et al., 1969). The Neethling vaccine was suggested to perform better in the control of LSD outbreaks than a SPPV based vaccine (Ben-Gera et al., 2015). The vaccine usually does not induce the febrile response and the development of skin lesions commonly associated with virulent LSDV infection. Occasionally, a mild vaccine-derived disease was reported (Ben-Gera et al., 2015; Katsoulos et al., 2018). Cattle inoculated with virulent and vaccine LSDV substantiated T-cells/macrophages response against LSDV infection (Ahmad et al., 2023; Fay et al., 2022; Haegeman et al., 2021; Kumar et al., 2024; Suwankitwat et al., 2023). Like other Poxviruses, LSDV possesses immunosuppressive strategies to counter host response (Liang et al., 2024), yet interaction of LSDV with cellular immunity is not fully understood.

Like many other Poxviruses (McFadden, 2005), LSDV has strong specificity to its host *in vivo* while showing wide promiscuity *in vitro*. LSDV can replicate in primary cultured cells and various cell lines originating from cattle (Binepal et al., 2001; Fay et al., 2020; Plowright and Witcomb, 1959; Weiss, 1968) as well as in cells from various non-permissive host animals including sheep (Babiuk et al., 2007; Plowright and Witcomb, 1959; Weiss, 1968), green monkey (Rhazi et al., 2021; Weiss, 1968), hamster (van Diepen et al., 2021; Weiss, 1968) and humans (Xie et al., 2024). *In vivo*, apart from dermal nodules, LSD may lead to extensive lesions in mucosal, pharyngeal, gastric membranes, and inflammation of internal organs of infected animals (Babiuk et al., 2008; Prozesky and Barnard, 1982; Sanz-Bernardo et al., 2021) suggesting wide tissue tropism of virulent LSDV. LSDV could also be isolated from white blood cell of infected cattle (Carn and Kitching, 1995; Menasherow et al., 2014).

Here, we characterized the cell tropism and replication of virulent and vaccine LSDV *in vitro*. We found that virulent and vaccine LSDV productively replicated with similar efficiencies in a bovine kidney cell line, in a bovine macrophage cell line and in primary bovine foreskin cells. In contrast, PBMCs failed to support significant viral DNA replication and release of infectious progeny, despite expression of early and late viral genes. We found that infectious LSDV suppressed the induction of ISGs’ RNA in PBMCs, which was observed when PBMCs were inoculated with heat-treated viruses. Despite failing to support LSDV replication, infected PBMCs transmitted the virus to recipient co-cultured MDBK cells and produced infectious foci, suggesting a potential role of PBMCs in viral dissemination.

## Materials and Methods

### Cell lines and viruses

Madin Darby kidney cells (MDBK) procured from ATCC (#CCL-2) were used for viral propagation and generation of recombinant LSDV strains. Bovine skin fibroblasts were prepared from a calf prepuce obtained from a local abattoir. Prepuce tissue was dipped briefly in a povidone-iodine solution, immersed in phosphate buffer saline (PBS), aseptically minced and incubated in a trypsin (0.25% w/v) EDTA (0.05% w/v) solution for 15 min. Tissue pieces were allowed to settle, supernatant containing detached cells were spun at 200 relative centrifugal force (RCF) for 5 min at room temperature, supernatant discarded and cells resuspended in culture media as described below and plated in T-25 tissue culture flasks. Foreskin cells were successfully propagated up to 25 passages with no apparent changes in proliferation. MDBK cells and skin fibroblasts were cultured in complete MEM media [minimal essential media (BI) supplemented with 10 % fetal bovine serum (v/v, BI), 1% penicillin-streptomycin-amphotericin B solution (BI) and 1% L-Glutamine (v/v, BI)] in a 5% CO2 humidified incubator at 37°C. Immortalized peritoneal bovine macrophage cells (BoMacs), a kind gift from Judith Stabel (USDA, (Stabel and Stabel, 1995)) , were cultured in complete RPMI media [RPMI-1640 media (BI) supplemented with 10% fetal bovine serum, 1% antibiotics and 1% L-Glutamine (v/v, BI)] in a 5% CO2 humidified incubator at 37°C.

### LSDV propagation

Virulent LSDV was isolated from a skin nodule of an infected cattle during the 2006 LSD outbreak in Israel (cell culture passage number-3). This LSDV field isolate and a Vaccine strain (OBP strain, batch no. 449, Onderstepoort Biological products SOC Ltd) were used in the described experiments and for genetic manipulation. These strains were propagated in MDBK cells as follows; MDBK cells were inoculated with the virus for 1 hour, washed with PBS and refreshed with complete MEM media. Cells were cultured for 8-10 days to allow a few rounds of viral replication and monitored daily for the development of a cytopathic effect (CPE) until the monolayer was near-completely disrupted. Supernatants were collected and subjected to three freeze and thaw cycles for the proper release of the cell-associated viruses. Supernatants were then centrifuged at 300 RCF for 5 min at room temperature to eliminate debris and stored in aliquots in -80 °C. Heat treatment (HT) of LSDV strains was done by incubating viral preparations at 55 °C for 30 minutes in a water bath.

### Viral titer determination

MDBK cells were used to titer LSDV. Tenfold dilutions of individual strains (at least 6 repeats for a single strain) were used to inoculate MDBK cells pre-seeded in 96 well plates. After 4-6 days, cells were inspected for the development of CPE and the endpoint dilution was recorded. TCID50 was calculated by the Reed and Muench method (Reed and Muench, 1938).

### Reagents

Minimal essential media, RPMI-1640 media, fetal bovine serum, penicillin-streptomycin-amphotericin B solution, L-Glutamine and DPBS were procured from Biological Industries (BI). Actinomycin D, carboxymethyl-cellulose (CMC), cycloheximide and phosphonoacetic acid (PAA) obtained from Sigma Aldrich. pGME T easy kit (Promega), Lipofectamine 2000 reagent (Invitrogen, USA), Gibson assembly cloning kit (NEB), Ficoll-paque plus solution (GE Healthcare), RNeasy kit (Qiagen), DNeasy kit (Qiagen), Dream Taq PCR master mix (Thermo Scientific), qPCR bio Sygreen blue mix (PCR biosystems), Turbo DNA free kit (Invitrogen), SensiFAST cDNA synthesis kit (Bioline), propidium iodide (PCR biosystems), BOVIGAM pokeweed mitogen (Applied biosystems).

### Construction of LSDV-GFP recombination plasmid

LSDV expressing GFP were created adopting previous strategies with some modifications (Chakrabarti et al., 1997; Hammond et al., 1997). In brief, a recombination plasmid was created from a linear pGME-T vector (Promega) by fusing it with a cassette having left and right flanking arms homologous with LSDV target sequences and a GFP sequence under a synthetic Vaccinia early/late promoter (Baur et al., 2010; Chakrabarti et al., 1997) (**Fig. S1**). The intergenic region between LSDV05 (left arm) and LSDV06 (right arm) having end-to-end orientation of ORFs was chosen as the genomic target of insertion in order to minimize possible interference with viral gene expression. Both flanking arms and the GFP coding sequence were amplified from viral DNA extracted from LSDV infected MDBK cells and pEGFPN-1 (Clontech) plasmid, respectively. PCR was performed using Phusion high fidelity DNA polymerase (NEB) with sets of primers introducing homology regions of LSDV genes and homology sequences to pGME-T for Gibson cloning (**Table S1**). Each fragment was separated through agarose gel electrophoresis and purified using Gene all Exspin kit (Genall Biotech Ltd). Purified PCR fragments and pGEM-T plasmid were joined by Gibson assembly cloning (NEB) as per the manufacturer’s instructions. Sanger sequencing was done to exclude any changes in the sequence of the GFP recombination cassette.

### Construction and purification of LSDV–GFP recombinant viruses

Confluent MDBK cells in 6-well plates were infected with LSDV field isolate or vaccine strains at MOI=0.5. After one hour of incubation, inoculum was removed and cells washed twice with PBS. Cells were replenished with 1 ml of MEM media containing 5% FBS. 0.3 ml Lipofectamine 2000 reagent (Invitrogen) complexed with 5 μg pGEM-T-LSDV-GFP plasmid was added to the wells gently. The culture plate was then placed in the incubator allowing viral replication and recombination events to produce GFP expressing LSDV. The next day, cells were gently washed with PBS and replenished with fresh complete MEM media. Cells were monitored up to 50 hours or until exhibiting CPE and the appearance of green-fluorescent foci. Media were then collected for plaque purification of GFP expressing LSDV recombinant viruses, conducted as follows: Media were inoculated to confluent MDBK cells for one hour, washed three times with PBS to remove the inoculum and overlaid with 1.5 ml of CMC solution (50% CMC (v:v), 10% MEM (v/v), 0.0025% Na2HCO3 (v/v), 10% Fetal bovine serum (v/v), 1% antibiotics, 1% L-glutamine (v/v) and 30% sterile water). The plate was then placed in the CO2 incubator. After 24-48 hours, one of the distinct fluorescent foci seen under the microscope was picked using a micropipette tip (avoiding non-fluorescent foci) and dipped in media for another round of purification. Three rounds of plaque purification were followed to get pure recombinant LSDV. The purity of recombinant LSDV from parental viruses was confirmed using PCR. DNA extracted from recombinant LSDV stocks (using DNeasy kit, Qiagen) was used for PCR with three sets of primers (**Fig. S1, Table S1**). PCR was performed using 2xDream Taq PCR master mix (Thermo Scientific) as per the manufacturer’s instructions, in brief 20 µl reaction was prepared with 10µl PCR mix (2x), 1µl of 10 µM forward and reverse primers, 3 µl template DNA and 5 µl distilled water. Thermal cycler program included: 95°C for 1 min, 35 cycles of: 95 °C for 15 sec, 58 °C for 30 sec and 72 °C for 85 sec, followed by terminal elongation at 72 °C for 10 min.

### Bovine peripheral blood mononuclear cells (PBMCs) isolation and culture

Peripheral blood was collected from the tail vein of Holstein calves in Becton Dickinson (BD) vacutainer heparin tubes. Calves bleeding for PBMCs preparation was approved by the Kimron Veterinary Institute Committee on Animal Research and Ethics (Ref. No. 102-2023). Blood cells were separated by density gradient centrifugation using a Ficoll-paque layer (density 1.086 g/ml, GE Healthcare,) with slight modifications to the manufacturer’s instructions. Blood was diluted in a 1:1 ratio in PBS (supplemented with 2% fetal bovine serum), layered on the top of Ficoll-paque in SepMate tubes (Stem cell technologies) and centrifuged at 1,200 RCF for 10 min at room temperature. In SepMate tubes, PBMCs float over red blood cells as a whitish layer. The PBMCs layer was decanted immediately into a new tube for re-centrifugation at 600 RCF for 8 min at room temperature. PBMCs were washed with PBS to pellet again at 600 RCF for 8 min at room temperature. Harvested PBMCs were suspended in complete RPMI medium. Viable cells were counted by trypan blue dye exclusion, seeded in culture plates and incubated overnight in a 5% CO2 humidified incubator at 37°C before using for LSDV infection.

### Flow cytometry analysis

MDBK, BoMac and bovine foreskin cells were collected using trypsinization. PBMCs were gently scraped with slow stirring in PBS. Collected cells were fixed for 10 min at room temperature with 4% neutral buffered formalin, passed through a 70-micron cell restrainer and processed in a FACS analyzer (BD FACS Calibur, Model E4856,). 20,000 events were recorded for every sample.

### Quantification of PBMCs viability

PBMCs viability was assayed by propidium iodide (PI) exclusion. Briefly, 10% PI (v:v) was added to uninfected or LSDV infected PBMCs for 10 min and cells were made up volume with PBS for FACS analysis. The percentage of viable cells was calculated as 100%- % PI positive cells.

### LSDV infection of PBMCs

PBMCs were seeded in 12-well plates at a density of 0.5 million cells per well. The next day, cells were inoculated with LSDV (recombinant or parental) strains, washed gently with PBS and supplemented with complete RPMI media. Where indicated, the viral DNA polymerase inhibitor PAA and the translation inhibitor CHX were added. Cells were lysed at designated time points to extract DNA/RNA for the quantitative analysis of viral DNA replication and gene expression. Where indicated PBMCs were stimulated by overnight incubation with pokeweed mitogen at a concentration of 5 µg/ml/million cells. The activity of pokeweed mitogen was confirmed by IFN-γ release to the medium of treated PBMCs as measured by a commercial IDscreen ruminant IFN-γ ELISA kit (IDvet).

### DNA /RNA extraction, Reverse transcription (RT), and quantitative PCR (qPCR)

DNA/ RNA extraction and purification from cells was conducted using the Qiagen RNeasy kit. Total RNA of all samples was quantified by using nanodrop reader (Thermo Scientific), DNA was removed using Turbo DNA free kit (Invitrogen) and then cDNA was synthesized using SensiFAST cDNA synthesis kit (Bioline) as per the manufacturer’s instructions. 10 µl qPCR reaction was composed of 5µl of 2x qPCR bio SyGreen blue mix, (PCR biosystems), 1µl forward primer (10µM), 1 µl reverse primer (10µM) and 3 µl diluted cDNA. qPCR reactions were run on a MIC real -time PCR cycler (Bio Molecular Systems, BMS) using the following program: pre denaturation at 95 °C- 3 min, 45 cycles of 95 °C- 15 sec, 55 °C- 15 sec and 60°C -25 sec. Samples without DNA digestions were used to check LSDV genome replication. Relative expression of viral or host genes were calculated using the delta-delta CT method (shown as a fold change in the histograms) using GAPDH as the reference gene.

### LSDV dissemination assay

PBMCs were infected with LSDV WT-GFP or LSDV Vac-GFP for two hours. After inoculation, PBMCs were washed once with PBS and then media was added to scrape cells from the well by a gentle stirring motion. PBMCs were then pelleted at∼ 600 RCF for 1 min, washed twice with PBS resuspended in MEM complete media and poured over pre seeded confluent MDBK cells. After 2 hours of incubation and settling of PBMCs (confirmed through microscopic observation), media was removed, and a 0.2% agarose overlay (in complete MEM media) was applied. Foci emergence was monitored by fluorescence microscopy and images were taken. After 48 hours, cells were fixed with 4% neutral buffered formalin and counter stained with Hoechst dye (1µg/ml). Plate imaging was performed using the Cytation microplate imager, version 5 (Biotek) using the GFP channel (Ex 470 nm/ Em 525 nm) for detection of fluorescent green cells foci and the DAPI channel (Ex 377 nm/ Em 454 nm) for detection of all cells (nuclear stained). Image analysis was conducted with the Cytation Gen5 software using a threshold circumference of 600µm ±50 for counting fluorescent cell foci.

### Fluorescence Microscopy

Mock infected**/**LSDV infected cells were imaged by Nikon TE 2000-U widefield microscope (Nikon, Japan) with a LED *p*E exciation system (CoolLED, UK) and a DS-Qi2 camera (Nikon, Japan) using NIS-Elements D imaging software (Nikon, Japan).

### Fluorometric determination of preformed GFP

LSDV/ LSDV-GFP stocks (prepared in MDBK cells) were quantified for GFP presence in the media by fluorometric approach using DeNovix DS-11 (Thermo fisher). In brief, 250 µl of media from LSDV GFP stocks were taken, excited at 488nm and measured emission at 506 nm. Media of uninfected cells / LSDV stocks used to check background fluorescence and normalization of RFU values.

### Statistical analysis

GraphPad Prism 9.1.0 software was used for statistical analysis and the creation of graphs. Data values are presented as mean± standard error of the mean (SEM). Statistical tests used to analyze experimental data are indicated in the figure legends.

## Results

### Generation of recombinant LSD viruses expressing GFP

In order to facilitate monitoring of LSDV infection and dissemination (by flow cytometry and by fluorescence microscopy), we have constructed recombinant versions of virulent and vaccine- attenuated LSDV expressing a GFP reporter gene (hereon referred to as LSDV WT-GFP and LSDV Vac-GFP, respectively). GFP was cloned under a Vaccinia virus synthetic early/late promoter (Baur et al. 2010) and targeted to the viral genome between LSDV05 and LSDV06 genes using homologous recombination without disrupting the flanking genes (**Fig. S1A**). MDBK cells were infected with either WT or Vac LSDV strains and transfected with a plasmid containing the recombination cassette. Recombinant viruses were isolated using three rounds of plaque purification from green-fluorescent foci of infected cells (**Fig. 1A**). The homogeneity of the isolated recombinant viruses was confirmed by PCR (**Fig. S1B**).

**Figure 1.**
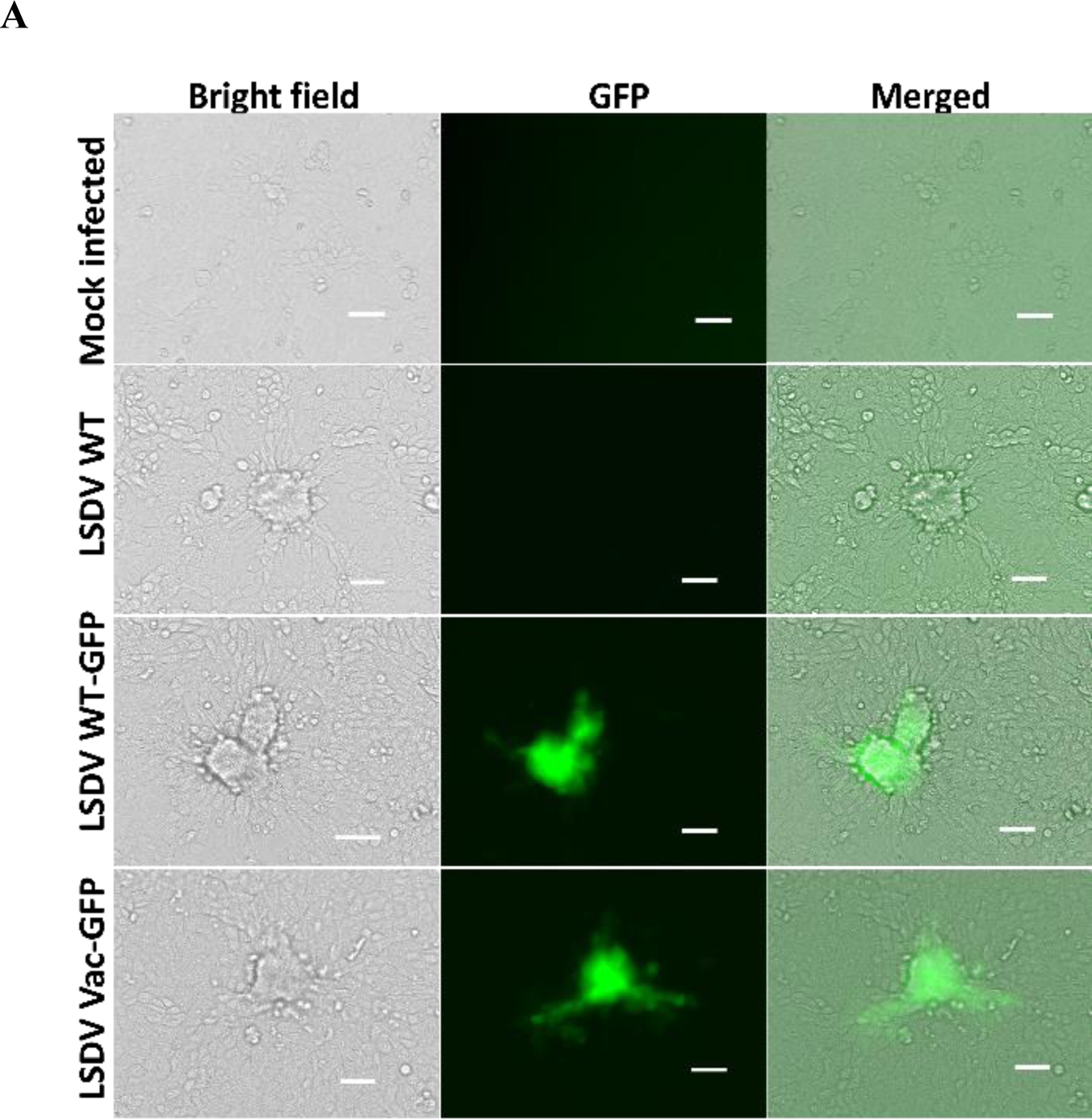
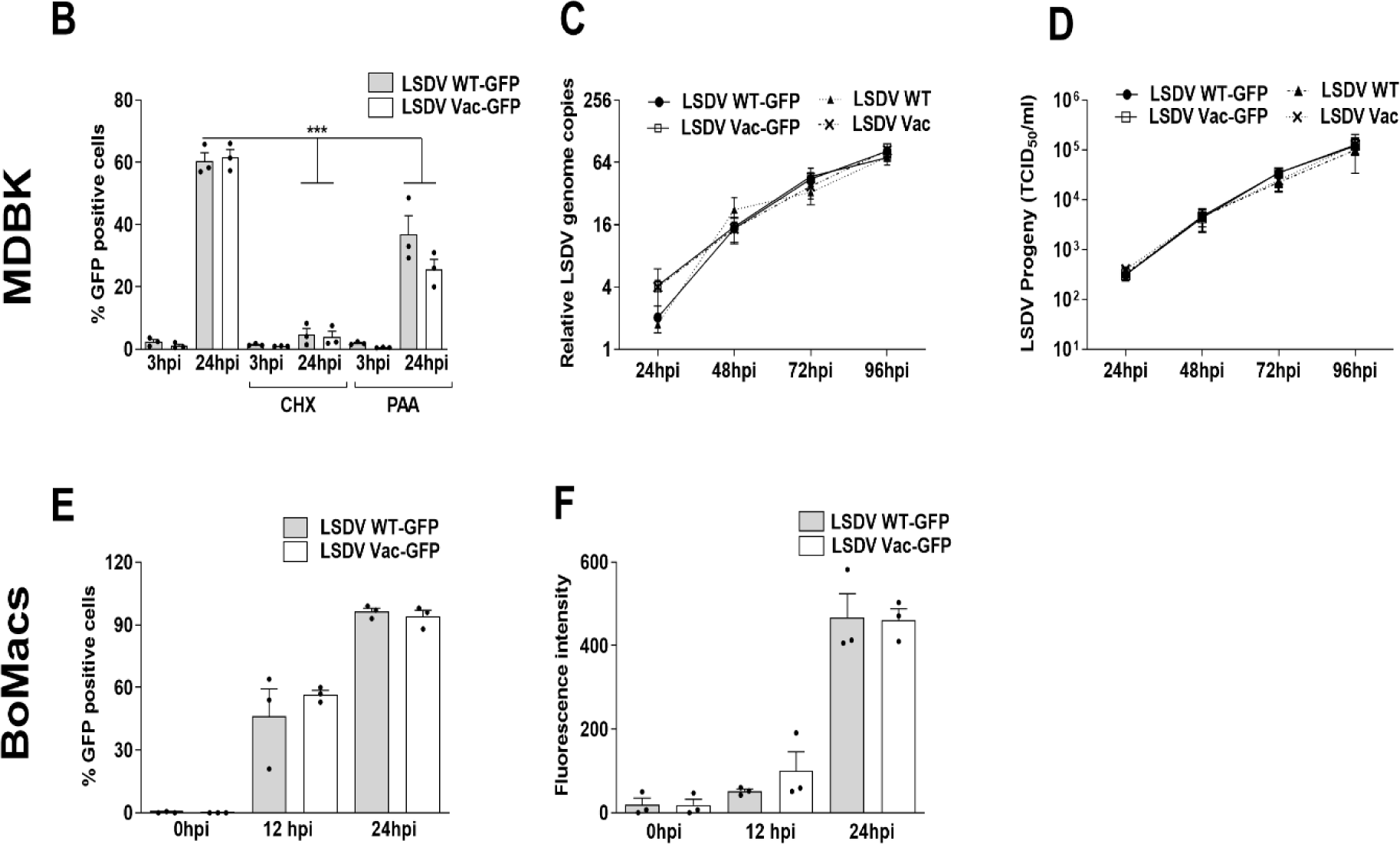
WT and Vaccine LSDV GFP recombinants efficiently replicate in MDBK and BoMac cells. **(A)** MDBK cells were inoculated with parental WT LSDV, Vac LSDV, plaque purified recombinant LSDV WT-GFP or LSDV Vac-GFP at MOI=3. After 48 h cells were imaged using epifluorescence microscopy to monitor emergence, morphology, and fluorescence of infection foci (Scale Bar 100µm). **(B)** MDBK cells were inoculated with LSDV WT-GFP and LSDV Vac-GFP at MOI=1, where indicated cells were pretreated with cycloheximide, (CHX, 20 µg/ml) or treated with the viral DNA polymerase inhibitor phosphonoacetic acid (PAA, 100 µg/ml). At the time points indicated cells were collected and the percentage of GFP positive cells was determined using flow cytometry. **(C-D)** MDBK cells were infected with either parental WT LSDV, Vac LSDV, LSDV WT-GFP or LSDV Vac-GFP at MOI=3 for 1h. At 1, 24, 48, 72 and 96hpi cells were harvested for viral DNA quantification (C) and infectious progeny (D). Relative LSDV DNA levels were determined by qPCR as fold change relative to 1hpi using GAPDH as an internal control (C). Infectious virus progeny was determined by TCID50 titration (D). **(E-F)** BoMac cells were infected with LSDV WT-GFP and LSDV VAC-GFP at MOI=3 for 1h (T=0). At the time points indicated, infected cells were collected and analyzed by flow cytometry. The percentage of GFP expressing cells (E) and the mean fluorescence intensity were determined (F). Data values are expressed as mean± SEM representing three biological replicates (B-F).

When MDBK cells were infected with LSDV WT-GFP or LSDV Vac-GFP, the percentage of GFP expressing cells (**Fig. 1B** and **Fig**. **S2A-C**) and the intensity of fluorescence signal per cell increased over time (**Fig. S2D**) corresponding with a progression of recombinant virus replication in host cells. GFP did not accumulate when LSDV-GFP infected MDBK cells were pre-treated with the translation inhibitor cycloheximide (CHX) (**Fig. 1B**) confirming that GFP is *de-novo* synthesized in infected cells. In addition, the percentage of GFP positive cells was markedly reduced when cells were treated with Phosphonoacetic acid (PAA, **Fig. 1B**), a known inhibitor of Herpes virus and Poxvirus DNA replication (Huang et al., 1976), supporting the dependence of GFP expression on viral replication. To determine whether the integration of the reporter gene has affected the replication properties of the LSDV strains, we compared the replication kinetics of the recombinant viruses in MDBK cells to that of their parental strains. MDBK cells were infected with parental or recombinant LSDVs at MOI=3. At 0, 24, 48, 72 and 96 hours post infection (hpi) cells were harvested for DNA extraction and viral DNA in cells was quantified using quantitative PCR (qPCR). Medium collected at the same time points was used to titer virus by TCID50. The kinetics of viral DNA replication (**Fig. 1C**) and production of infectious virus progeny (**Fig. 1D**) were similar between parental and GFP recombinant LSDVs confirming that integration of GFP did not impede viral replication. Replication kinetics were also similar between WT and Vac LSDVs.

*In vivo*, LSDV is found in highest titers and is most easily isolated from skin nodules of infected cattle (Davies et al., 1971; Moller et al., 2019; Tuppurainen et al., 2005). Therefore, we chose primary cells of skin source (fibroblast cells derived from prepuce of a calf) to further characterize LSDV recombinants. Like MDBK cells, primary fibroblasts infected with LSDV WT-GFP or LSDV Vac-GFP gave rise to increasing GFP fluorescence over time (**Fig. S2 E-H**).

### LSDV WT-GFP and LSDV Vac-GFP productively replicate in a bovine macrophage cell line

Cattle infected with virulent LSDV show a different immune response than ones infected with the attenuated vaccine strain, marked by a strong inflammatory response of the former (Haegeman et al., 2021; Moller et al., 2019). We thus assessed whether WT and Vac LSDV differ in their ability to infect and replicate in cells of the immune system. Previous histopathological and immunohistochemical analysis of tissues from LSDV infected cattle detected typical Poxvirus inclusion bodies (Prozesky and Barnard, 1982) and LSDV antigens (Awadin et al., 2011; Babiuk et al., 2008) within macrophages in the skin and in lymph nodes. We found that both virulent and attenuated LSDV could infect and replicate in immortalized peritoneal bovine macrophage cells (BoMac cells, (Stabel and Stabel, 1995)) with similar efficiencies (**Fig.1E-F**). These transformed monocytic (BoMac) cells have lost some of their functional properties (Sager et al., 1999). We therefore tested LSDV replication and immune response in primary peripheral white blood cells.

### Primary white blood cells are susceptible to LSDV infection yet do not support significant viral replication

Preparations of buffy coats from blood of cattle either naturally or experimentally infected with LSDV were shown to contain viral DNA, as well as infectious virus (Carn and Kitching, 1995; Menasherow et al., 2014; Sanz-Bernardo et al., 2021). However, active viral replication in these cells was not confirmed. We have isolated PBMCs from naïve calves (neither previously exposed to LSDV nor vaccinated against the disease). After seeding in culture plates and overnight incubation, cells were infected with LSDV WT-GFP or LSDV Vac-GFP at MOI=3 (this MOI evaluation was based on titration of stocks on MDBK cells). After 2h of inoculation (T=0) ∼ 5% of PBMCs became GFP positive, increasing after 48 hours to an average of ∼25% of cells (i.e. effective MOI in PBMCs=0.25) suggesting lower susceptibility to infection of PBMCS compared to MDBK cells (**Fig. 2A**). Of note, we observed significantly higher variation in the percentages of GFP positive cells with the different independent preparations of PBMCs (**Fig. 2A)**, compared to the low variation found when infecting the homogenous populations of bovine cell lines or primary fibroblasts (**Fig. 1B, E and Fig. S2C, G)**. LSDV WT-GFP and LSDV Vac-GFP were equally infectious to adherent PBMCs (**Fig. 2A**). When inspecting the intensity of the GFP signal in infected PBMCs it was not significantly increased over the course of infection (**Fig. S3A**) unlike the dramatic increase seen in infected MDBK, fibroblasts and BoMac cells (**Fig. S2D, Fig. S2H** and **Fig. 1F** respectively). Moreover, viral DNA copies in LSDV infected PBMCs had not significantly increased (**Fig. 2B, left bars**) and media collected from infected PBMCs indicated only a minor increase (*ca* 3-fold) in viral DNA (**Fig. 2B, right bars**) or in viral infectious progeny (**Fig. 2C**) during the course of PBMC infection with either LSDV WT-GFP or LSDV Vac-GFP. We further monitored viral replication in pokeweed mitogen stimulated PBMCs, since cellular proliferation might support viral replication, as shown for other viruses (van der Muelen et al 2001, Rey and Kenneth 1996). However, we found only a minor increase in LSDV DNA copies during the course of infection of stimulated PBMCs similar to that found in non-stimulated cells (**Fig. S3B**).

**Figure 2.**
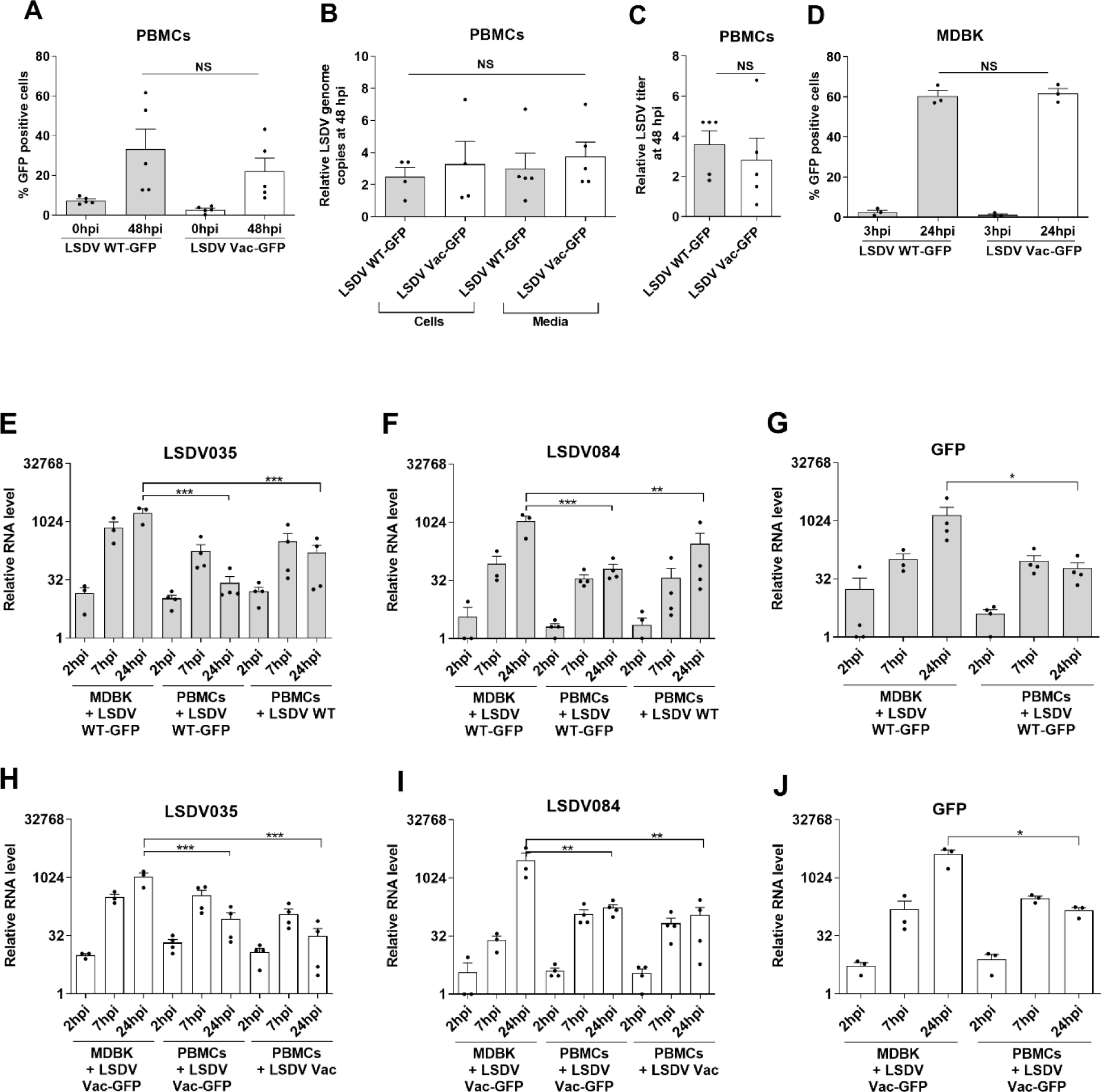
PBMCs are susceptible to LSDV-GFP (WT and Vac) infections but poorly support viral replication in contrast to MDBK cells. (A-C) Naïve PBMCs were inoculated with LSDV-GFP strains at MOI=3 for 2 hours (T=0) and further incubated for 48h. At both time points, cells were collected for flow cytometry analysis (A). DNA extracted from infected PBMCs (B, left bars) and collected media (B, right bars) were assayed for relative genome copy versus time point 0hpi. Medium was also assessed for the relative increase in virus progeny titer (TCID50) (C). Titration involved half log dilution to allow better resolution of titers. **(D)** MDBK cells a were inoculated with LSDV-GFP strains in parallel to PBMCs as described above yet at MOI=1 and analyzed for the percentage of GFP positive cells by flow cytometry. **(E-J)** MDBK cells or PBMCs were inoculated with LSDV WT-GFP, LSDV WT (E-G), LSDV Vac- GFP or LSDV Vac (H-J) as indicated, at MOI=1 for 1 hour. After incubation, cells were washed and replenished with fresh media. At 1h, 2h, 7h and 24h, cells were lysed, and nucleic acids extracted. Relative changes in the levels of transcripts encoding the reporter gene GFP and viral genes (LSDV035 and LSDV84) were determined by RT-qPCR, normalized to GAPDH RNA (internal control) and fold change calculated against T=0 (i.e. 1 hour inoculation). Three independent sets used to plot the figures and data values presented as mean± SEM. One way ANOVA followed with Tukey’s post-hoc test used to compare all groups (E-F, H-I). Ratio paired t- test was used in G and J. ∗p < 0.05, ∗∗p < 0.01, ∗∗∗p < 0.001, NS- non-significant.

To explore the possibility that the GFP signal in infected PBMCs represents carryover of preformed GFP from LSDV-GFP viral stocks (produced in MDBK cells) rather than infection induced expression of GFP in PBMC, we have conducted two control experiments: 1) We have inoculated PBMC with LSDV WT-GFP and LSDV Vac-GFP which were heat treated (HT) at 55°C for 30 min causing their titers to be reduced by 100 fold (**Fig. S4 C**). PBMCs inoculated with HT GFP viruses did not show green fluorescence unlike cells inoculated with fully infectious viruses (**Fig. S4A**). Importantly, GFP has high thermal stability (Bokman and Ward, 1981) and we have confirmed that heat treatment had not affected the fluorescence of preformed GFP, which may have been found in LSDV GFP stocks (**Fig S4D**). 2) In a second control experiment, treating PBMCs with the translation inhibitor CHX prior to infection with LSDV-GFP viruses prevented the appearance of green fluorescence in cells (**Fig S4B**). Taken together, our results show that GFP is *de-novo* synthesized in LSDV-GFP infected PBMCs and is indicative of the susceptibility of PBMCs to LSDV WT-GFP and LSDV Vac-GFP infection. We asked whether failure of PBMCs to productively support LSDV replication is associated with virus induced cell death. However, we did not find any significant difference in viability between uninfected PBMCs and PBMCs infected with either LSDV WT-GFP or LSDV Vac-GFP as determined by PI exclusion (>90% viability for all the tested sets, **Fig S5**).

### LSDV exhibited progressive viral genes expression in infected PBMCs

We further investigated whether LSDV infection of PBMCs is associated with a failure to execute its transcriptional program. We picked four LSDV genes (LSDV035, LSDV076, LSDV084 and LSDV089) to follow viral transcription in infected cells. These viral genes were chosen on the criteria of homology to Vaccinia virus (Tulman et al., 2001) to represent early and late viral genes (**Table S1)** . We used the Vaccinia virus expression annotation, as the knowledge on the regulation of LSDV gene expression is very limited (Fick and Viljoen, 1994). We first characterized the expression of these viral genes and the reporter GFP gene during productive LSDV-GFP replication in MDBK cells and fibroblasts (**Fig. 2E--J, Fig. S7A-I**). All four LSDV genes and GFP increased in abundance as early as 2 hpi and reached to a similar fold increase in abundance at 24 hpi. LSDV035 (**Fig. 2E, H and Fig. S7E**) and LSDV076 (**Fig. S7A, C and F**) were relatively more abundant than the other LSDV genes and GFP at early time points (2 hpi and 7 hpi) in both MDBK cells and fibroblasts. The dependency of late viral gene expression on viral DNA replication was previously demonstrated for Vaccinia virus (Keck et al., 1990; Yang et al., 2011). The kinetics of LSDV gene expression in MDBK cells revealed that LSDV084 gene met this criterion of a late gene since it was inhibited by PAA treatment (that interfered with viral genome replication, **Fig. S6A**) while LSDV035 remained unaffected by PAA supporting its classification as an early LSDV gene (**Fig. S6B**). The expression pattern of all tested viral genes was not significantly different between WT and Vac LSDV. MDBK cells infected with HT LSDV have shown lower induction of viral genes and GFP (**Fig. S8**). The lack of complete inhibition of viral transcription may be due to incomplete inactivation of the viruses with the heat procedure, that we have used (**Fig. S4C**).

We then followed the expression of LSDV genes in infected PBMCs. We found that RNA of all the selected viral genes (LSDV035, 076, 084, 089) and GFP increased in abundance in PMBCs at 7 hpi and at 24 hpi (**Fig. 2E-J and Fig. S7A-D**). Viral and GFP RNA levels were significantly reduced when HT viruses were used for infecting PBMCs (**Fig. S9**). LSDV infected PBMCs showed only a mild decrease in the levels of viral transcripts in response to protein synthesis arrest by CHX treatment compared to untreated cells, suggesting that (at least up to 24 hpi) their transcription is driven by preformed proteins (**Fig. S10**). Of note, the levels of viral gene induction in LSDV-GFP infected PBMCs (∼100-fold increase at 24 hpi, **Fig. 2E-J**) was significantly lower than that achieved in MDBK cells (>1000-fold at 24hpi, **Fig. 2E-J**). PBMCs infected with parental (non-recombinant) LSDV has shown a similar expression pattern as LSDV-GFP viruses (**Fig. 2E- J and Fig. S7A-D**).

### Infectious LSDV suppressed the induction of Interferon Stimulated Genes (ISGs) in PBMCs

It was previously shown that an intact type I interferon response may serve as a barrier for Poxvirus productive replication (Wang et al., 2004). We wished to determine if the nonproductive LSDV replication in PBMCs is associated with a cellular anti-viral response. To this end, we followed the kinetics of expression of ISGs IFIT1, IFIT2, IFIT3, ISG15 and IFITM3 in PBMCs inoculated with virulent and attenuated LSDV and their HT preparations (**Fig. 3A-E**). All tested ISGs were significantly induced by the HT viruses but not by infectious LSDV. IFITM3 RNA levels were less affected by HT viruses showing ca 10-fold increase while other tested ISGs were induced ca 100-fold. Of note, IFIT1-3 were induced to higher levels by HT Vac-LSDV compared to HT WT- LSDV. In MDBK cells, which support productive replication of LSDV, neither infectious nor HT viruses significantly induced the levels of ISGs’ RNAs **Fig. S11**).

**Figure 3.**
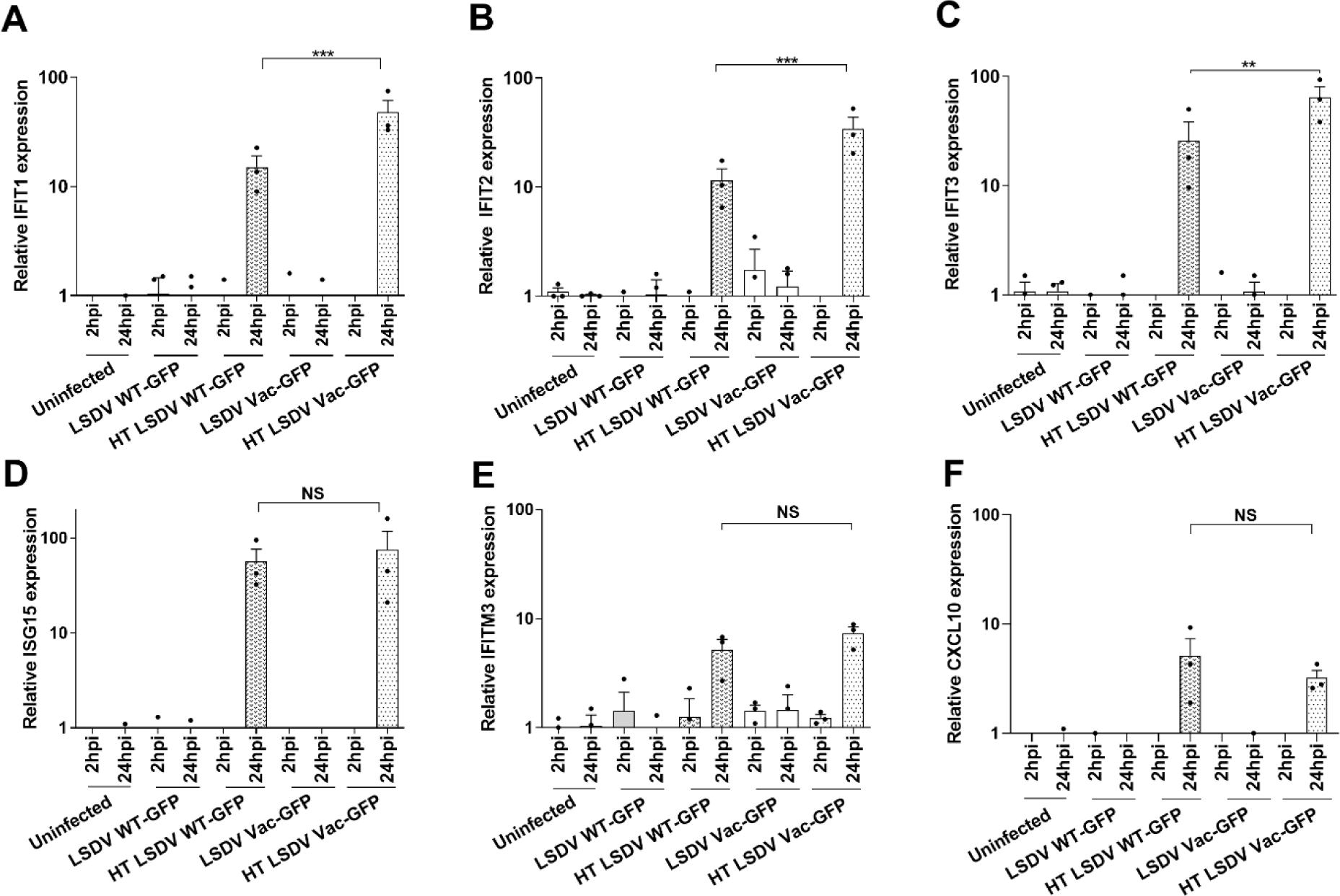
Heat-treated but not infectious LSDV strains upregulate ISGs and CXCl10 transcripts in PBMCs. (A-F) A comparative analysis of ISGs (IFIT1,2,3, ISG15 and IFITM3) expression performed in uninfected or PBMCs inoculated with infectious LSDV-GFP at MOI=1 or with equal volumes of HT LSDV-GFP viruses for 1h. After wash (T=0) and at the time points indicated, cells were lysed, and nucleic acids extracted. Relative changes in the levels of transcripts encoding ISGs were monitored by RT-qPCR, normalized to GAPDH RNA (internal control) and fold change calculated against T=0 (i.e. 1 hour incubation of uninfected cells). The relative expression of viral genes for this experiment is shown in **Fig. S9**. Fold changes were plotted in bar graphs as mean± SEM for three independent biological sets. One way ANOVA was used to compare all means and post-hoc Tukey’s test followed to test significance levels. ∗∗p < 0.01, ∗∗∗p < 0.001, NS- non-significant.

IFNγ is a key immune stimulatory cytokine secreted by various immune cells in response to viral infection (Billiau and Matthys, 2009). We found that HT LSDV but not infectious viruses induced the expression of the IFNγ induced chemokine gene CXCL10 **(Fig. 3F**) in PBMCs at 24 hpi, with similar induction achieved by HT WT and HT Vac LSDV. Taken together, we suggest that LSDV suppresses both type I and Type II interferon responses.

### Infected PBMCs can disseminate LSDV to neighboring permissive cells

We found that LSDV could infect adherent PBMCs, yet its replication in these cells appeared to be negligible. We wished to determine whether LSDV infected PBMCs may transmit the virus to neighboring susceptible cells. To address viral dissemination, PBMCs were first infected with WT or Vac LSDV-GFP for 1 hour, extensively washed and seeded on a monolayer of MDBK cells which were then overlaid with semi-solid media and incubated at 37°C for 48h. Single green fluorescent MDBK cells were apparent as soon as 12 hours of co-culturing with infected PBMCs and discrete fluorescent cell foci were formed by 24 hours (data not shown), suggesting transfer of infectious LSDV from PBMCs to MDBK cells. Fluorescent foci in the co-cultured MDBK cells increased in size from 24h to 48h (**Fig. 4A**). We did not find any significant difference in the ability of WT and Vac LSDV-GFP strains to disseminate to MDBK cells as apparent from foci counts (**Fig. 4B**). Altogether, both WT and Vac LSDV poorly replicate in PBMCs but may use these cells to disseminate to neighboring cells.

**Figure 4.**
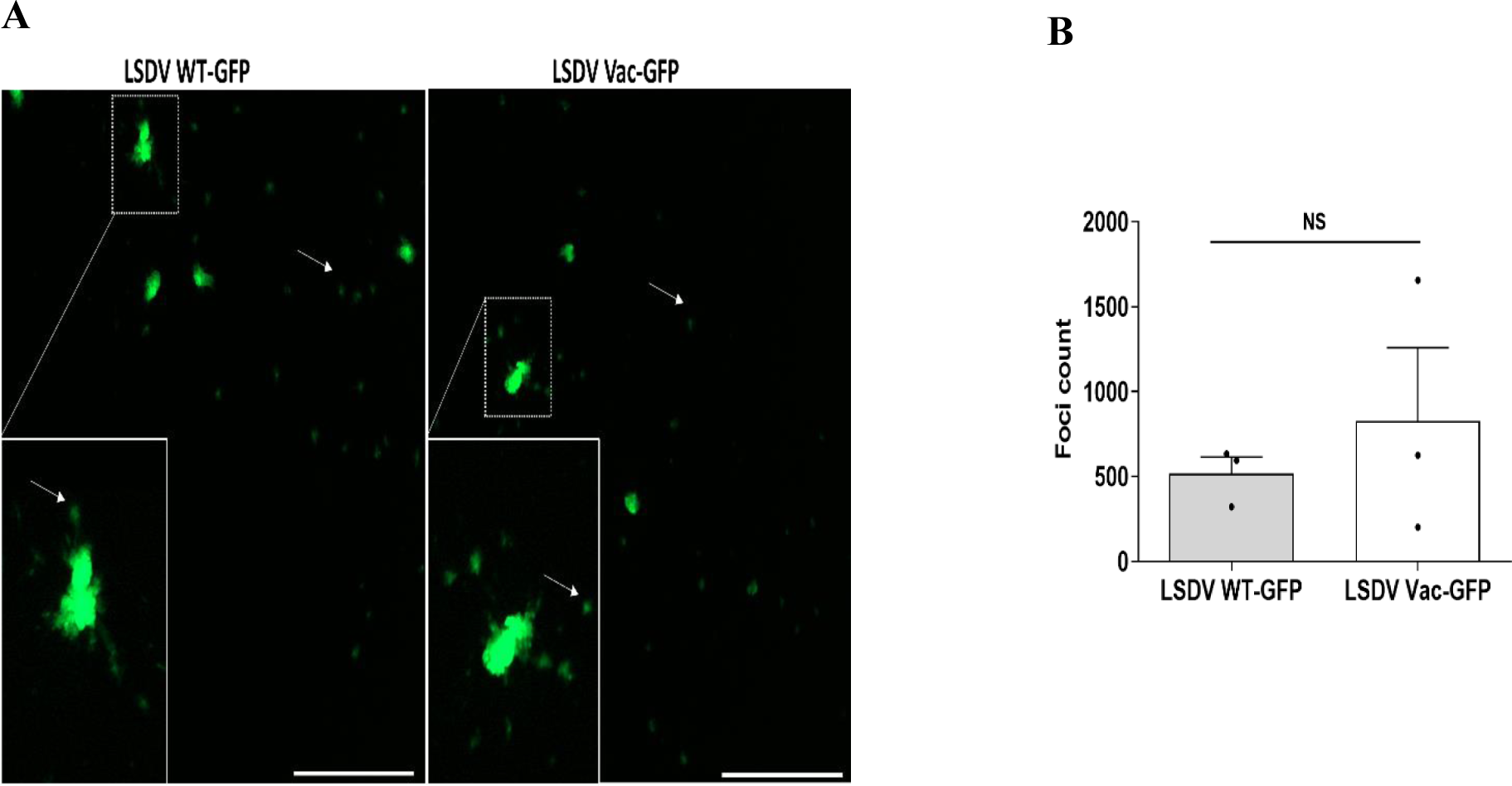
WT and Vac LSDV GFP viruses disseminated to MDBK cells from infected PBMCs. (A-B) PBMCs were infected with WT and Vac LSDV-GFP at MOI=1 for 2 hours, extensively washed, collected, seeded on a monolayer of MDBK cells, overlaid with semi solid media and incubated at 37°C for 48h. After formalin fixation, plates were scanned using the Cytation microplate reader version 5 (Agilent Biotek). Fluorescent foci (A, representative images, scale bar 1000 µm) were counted using the Cytation imaging software (B). Three independent repeats were used to plot the histogram and values were presented as mean± SEM. Paired t-test was used for statistical analysis. NS- non- significant.

## Discussion

*In vivo*, LSDV accumulates to highest loads in skin lesions of infected cattle yet viral physical particles, viral DNA and infectious viruses were found in various tissues and organs as well as in whole blood (Babiuk et al., 2008; Moller et al., 2019). Fever, ocular and nasal discharge and the development of skin nodules are typically associated with virulent LSDV infection. These clinical signs are rare and milder, when apparent in cattle vaccinated with the attenuated Neethling strain (Ben-Gera et al., 2015; Katsoulos et al., 2018). The mechanism behind this marked phenotypic difference is not well understood. A possible explanation is a differential effect of virulent and attenuated LSDV on the host innate and acquired immune systems. Pathogenesis of virulent and attenuated LSDV may also involve differences in cell and tissue tropism, in replication and dissemination properties of these viruses. We have characterized the replication of LSDV in bovine cultured cells using recombinant GFP reporter versions of virulent (WT) and vaccine- attenuated viruses (**Fig. 1, Fig. S1**). We exploited the ability of Pox viruses s to incorporate inserts into their genome by homologous recombination, as previously used to create recombinant LSDV based vaccines for rabies, bovine ephemeral fever, and Rift valley fever (Aspden et al., 2003; Aspden et al., 2002; Douglass et al., 2021; Wallace and Viljoen, 2005). LSDV WT-GFP and LSDV Vac-GFP replicated in MDBK cells with similar kinetics compared to their parental strains and to each other (**Fig. 1C-D**), in agreement with previously shown efficient isolation and propagation of both LSDV types in these cells (Fay et al., 2020). LSDV WT-GFP and LSDV Vac-GFP viruses also replicated with similar efficiencies in primary bovine foreskin cells (**Fig. S2E-H**). *In vivo,* skin nodules do not form in most vaccinated cows, and the ones formed in a minority of LSD vaccinated cows are smaller and contain less virus than those found in cattle infected with virulent virus (Ben-Gera et al., 2015; Katsoulos et al., 2018). Our *in vitro* findings suggest that this phenotypic difference is not due to reduced tropism of attenuated LSDV to skin cells. Similarly, we found no differences in replication of LSDV WT-GFP and LSDV Vac-GFP in immortalized peritoneal bovine macrophage cells (**Fig. 1E-F**), while *in vivo* only the virulent LSDV strain was widely detected in lymph nodes and in whole blood where macrophages may be found (Moller et al., 2019).

LSDV may encounter circulating bovine blood cells upon introduction through biting insects. Interestingly, under experimental conditions, intravenous injection of the virus causes a more reproducible and disseminated disease compared to intradermal inoculation (Carn and Kitching, 1995). Preparations of buffy coats from blood of naturally and experimentally infected cattle contain LSDV DNA as well as infectious virus (Babiuk et al., 2008; Carn and Kitching, 1995; Menasherow et al., 2014; Moller et al., 2019; Sanz-Bernardo et al., 2021). The detection of LSDV in cattle’s blood usually coincides with an inflammatory response (Babiuk et al., 2008; Moller et al., 2019; Sanz-Bernardo et al., 2021; Tuppurainen et al., 2005). Viral loads in the blood correlated with the severity of clinical signs in infected animals (Moller et al., 2019; Sanz-Bernardo et al., 2021). Under experimental mechanical transmission of LSDV by biting insects, LSDV DNA was detected in the blood of recipient cattle prior to or coinciding with the appearance of skin nodules, yet viremia peaked only several days later (Issimov et al., 2020; Paslaru et al., 2021; Sohier et al., 2019). Nonetheless, it was unclear if LSDV actively replicates in blood cells. We have found that adherent PBMCs isolated from naïve cattle are susceptible to infection with LSDV WT-GFP and LSDV Vac-GFP, as marked by GFP appearance (**Fig. 2A**). The GFP signal in PBMCs infected with reporter viruses was a result of virus driven *de-novo* synthesis of GFP, as accumulation of fluorescence was blocked by pre-treating cells with a translation inhibitor or by heat treatment of the inoculating viruses (**Fig. S4**). However, PBMCs failed to support productive LSDV replication as judged by no significant increase in viral DNA in the cells and negligible secretion of infectious progeny into the medium (**Fig. 2B-C**). The limited ability of LSDV to replicate in PBMCs was not correlated with virus induced cell death (**Fig. S5**) and was not enhanced by PBMC stimulation with pokeweed mitogen (**Fig. S3B**).

The regulation on Poxvirus transcription was extensively studied using Vaccinia virus (Moss, 1990). Upon entry, Vaccinia virus undergoes initial uncoating forming viral cores within which early viral genes are transcribed. After a lag period, a second uncoating occurs followed by viral DNA replication and transcription of late viral genes. Inhibitors of translation prevent uncoating of viral cores leading to prolonged and enhanced transcription of early viral genes (Moss, 1990). We found that both early (LSDV035 and LSDV076) and late (LSDV084, LSDV089) viral genes were transcribed in LSDV infected PBMCs (**Fig. 2E-J, Fig. S7 A-D**). CHX treatment of PBMCs did not inhibit LSDV transcription (**Fig. S10**) in line with progression of early viral gene expression using preformed viral proteins. Heat treatment of LSDV caused a pronounced inhibition of viral RNA accumulation in infected cells (**Fig. S9**) in agreement with its documented effect on rabbit poxvirus transcription (Kates and McAuslan, 1967). Taken together, LSDV transcription occurs in infected PBMCs but does not lead to efficient viral DNA replication and to completion of the viral replication cycle. The mechanism behind LSDV abortive infection of PBMCs remains to be elucidated. We have not observed significant differences in viral transcripts levels between WT and Vaccine LSDV in either MDBK cells, fibroblasts or in PBMCs (**Fig. 2E- J, Fig. S7**). Interestingly, ovine PBMCs inoculated with WT Sheepox virus resulted in higher viral RNA levels compared to those found in PBMCs infected with an attenuated vaccine strain. However, whether either of these viruses completed their replication cycle in PBMCs to produce infectious progeny was not shown (Chibssa et al., 2021).

Unlike many other viruses, post entry events rather than the availability of receptors restrict Poxvirus cellular tropism (McFadden, 2005). Subsets of 3T3 murine fibroblasts which were non-permissive to Myxoma virus replication, showed comparable levels of viral binding, entry, and early gene expression to permissive subsets 3T3 cells (Johnston et al., 2003). Similarly, Vaccinia virus equally associated with a fibroblast cell line and with monocyte-derived dendritic cells (DCs). However, while fibroblasts supported viral replication and late gene promoter activation, infection of DCs was abortive and allowed activation of early, but not late viral promoters (Jenne et al., 2000). Likewise, human primary blood monocytes differentiated into macrophages with either human AB serum, granulocyte macrophage colony-stimulating factor (GM-CSF) or macrophage colony-stimulating factor (M-CSF) equally allowed Vaccinia virus binding and early viral gene expression while only the latter two supported productive viral replication (Byrd et al., 2014). In fact, the ability of Poxviruses to infect and express early viral genes in human cells without completing a productive infectious cycle may be advantageous when using such viruses as platforms for vaccines or for genetic and anti-cancer therapies.

We sought to find if the nonproductive LSDV infection of PBMCs is associated with an interferon induced antiviral response. We found that neither WT nor Vaccine LSDV induced the expression of ISGs in PBMCs. In contrast, ISG were strongly induced in PBMCs inoculated with heat treated viruses suggesting that by an unknown mechanism infectious LSDV suppresses the IFN response in PBMCs (**Fig. 3)**. It also suggests that a certain component of the inactivated viral particle is causing PBMCs to mount an IFN response. Similarly, it was previously shown that heat-treated but not infectious Vaccinia virus induced the secretion of IFNα from plasmacytoid dendritic cells and was dependent on endosomal acidification and TLR7 signaling (Cao et al., 2012). Interestingly, HT Vaccine LSDV was more potent than the HT WT virus in inducing IFIT1-3 RNAs in PBMCs. Whether this results from a quantitative difference (amount of physical particles) or a qualitative difference (particle composition) between Vaccine and WT preparations remains to be studied. Our findings suggest that vaccination with a partially inactivated LSDV vaccine strain may exhibit a beneficial IFN stimulatory response, which would be suppressed in case of vaccination with a live attenuated vaccine. Such partially inactivated vaccine formulation may be advantageous by combining IFN stimulation (driven by dominance of inactivated particles) with the capacity to raise cell-mediated immunity (driven by residual infectivity).

It was recently shown that infectious but not UV inactivated LSDV induced the expression of IFNꞵ RNA in MDBK cells at 36 hpi and 48 hpi but not at earlier time points (Liang et al., 2024). Three ISGs (ISG54, ISG56 and Mx1) were also induced by infectious LSDV starting from 24 hpi (ca 2- fold) and increasing at 36 hpi and 48 hpi. In our experiments, ISGs were not significantly changed in MDBK cells infected with either infectious or HT LSDV (Fig. S11). This may reflect differences in the kinetics or potency of inducing IFN by the different field isolates used in Liang and coworker’s study and in our experiments.

Viruses of the Pox family are known for their ability to disseminate through cell-to-cell spread (Roberts and Smith, 2008). In order to evaluate a potential role of PBMCs in LSDV dissemination, we have conducted co-culture experiments. We found that despite failing to productively support viral replication, LSDV infected PBMCs transmitted the virus to co-cultured permissive MDBK cells (**Fig. 4**). This dissemination may have been carried out by infectious LSDV particles, which engaged PBMCs externally without entering these cells, or otherwise by LSDV transferred from PBMCs after entry, or by both of these routes. Cell-to-cell transmission of replicative naked viral genomes is also possible (Cifuentes-Munoz et al., 2020). Our findings suggest a possible role of PBMCs as carriers of LSDV within the host, which may disseminate the virus to multiple permissive cells and tissues. Human monocyte-derived macrophages which supported productive replication of Vaccinia virus, have formed virion-associated structures which may contribute to cell-cell spread (Byrd et al., 2014). Moreover, white blood cells were also demonstrated to serve as potential virus carriers without supporting viral replication. For example, foot and mouth disease virus (FMDV), an important pathogen of cattle and other livestock animals, was shown to be phagocytosed by macrophages and remained infectious within these cells despite showing no viral RNA synthesis. FMDV infected macrophages were also shown to transfer the virus to co-cultured permissive BHK-12 cells (Rigden et al., 2002). The mechanism of LSDV cell-to-cell transmission remains to be elucidated.

## Conclusions

We have found that virulent and attenuated LSDV efficiently replicate in MDBK, primary fibroblasts and BoMac cells, while poorly replicating in primary white blood cells. White blood cells can nonetheless support LSDV dissemination to permissive cells. Whether *in vivo* a subset of blood cells not cultured under our experimental conditions may support efficient replication of LSDV is currently unknown. Our findings support suppression of ISGs expression in PBMCs by infectious LSDV. The mechanisms limiting LSDV replication in PMBC are yet to be discovered and may shed light on the intricate interactions of virus and host cells.

## Supporting information

Supplemental data for Kumar et al

## Acknowledgements

We thank Judith Stabel (USDA) for kindly providing the BoMac cell line. We thank Avi Eldar and Ezra Rozenblut (RIP) (Kimron Veterinary Institute) for their assistance in isolation and culturing of primary bovine fibroblasts. The help of Kimron Veterinary staff - Boris Leibovich and Daniel Landau in providing cattle’s blood is highly appreciated. Assistance by Ahuva Friedman, Saveliy Kirillov and Maria Billan (Rouvinski’s lab, HUJI) with Cytation plate reader acquisition and analysis is highly acknowledged.

## Conflict of interest

All authors declare no conflict of interest.

## Funding sources

Israel Ministry of Agriculture Grant 33-08-0007 to Dr Sharon Karniely (Kimron Veterinary Institute, Bet Dagan, Ministry of Agriculture, Israel) and Dr Alexander Rouvinski (Hebrew University of Jerusalem, Jerusalem, Israel).

## Notes

### Competing Interest Statement

The authors have declared no competing interest.

