## Supplemental data for Kumar et al for "Primary bovine white blood cells support dissemination of Lumpy Skin Disease Virus while suppressing viral replication"

### Supplementary data and figures

**A**

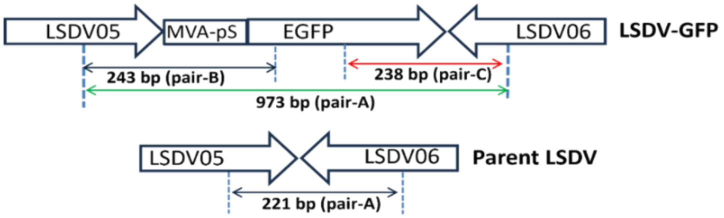


**B**

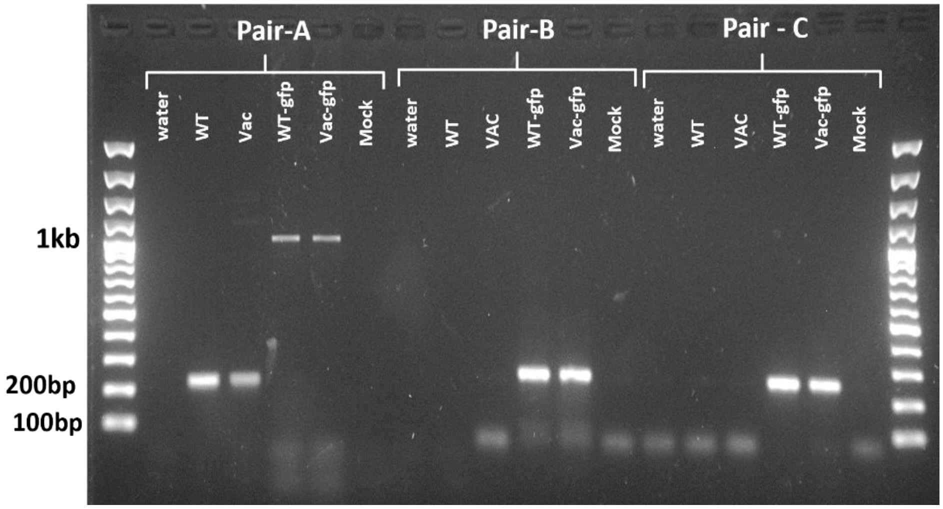


**Figure S1. Generation of recombinant LSDV viruses expressing GFP.**

**(A)** GFP was cloned under a Vaccinia virus synthetic promoter (MVA-pS, (Chakrabarti et al., 1997)) flanked by sequences with homology to the ends of LSDV05 and LSDV06 genes. Wide arrows in the scheme represent the orientations of viral ORFs. Thin colored arrows represent predicted amplicon sizes using the specified primer sets (detailed in **Table S1**).

**(B)** LSDV-GFP viruses were screened for homogeneity by endpoint PCR. DNA extracted from MDBK cells either mock infected or infected with parental and recombinant LSDV strains were amplified by PCR using three primer sets designated as pair A, B and C: Pair A- left flanking arm (LSDV05) + right flanking arm (LSDV06); Pair B- left flanking arm (LSDV05) + mid EGFP sequence and Pair C- mid EGFP sequence + right flanking arm (LSDV06).Amplicons were then separated by agarose gel electrophoresis, Amplification of DNA from recombinant viruses with flanking primers gave rise to a single amplification product corresponding to the target viral genomic site including integration of the EGFP cassette with no traces of an amplicon derived from parental LSDV DNA.

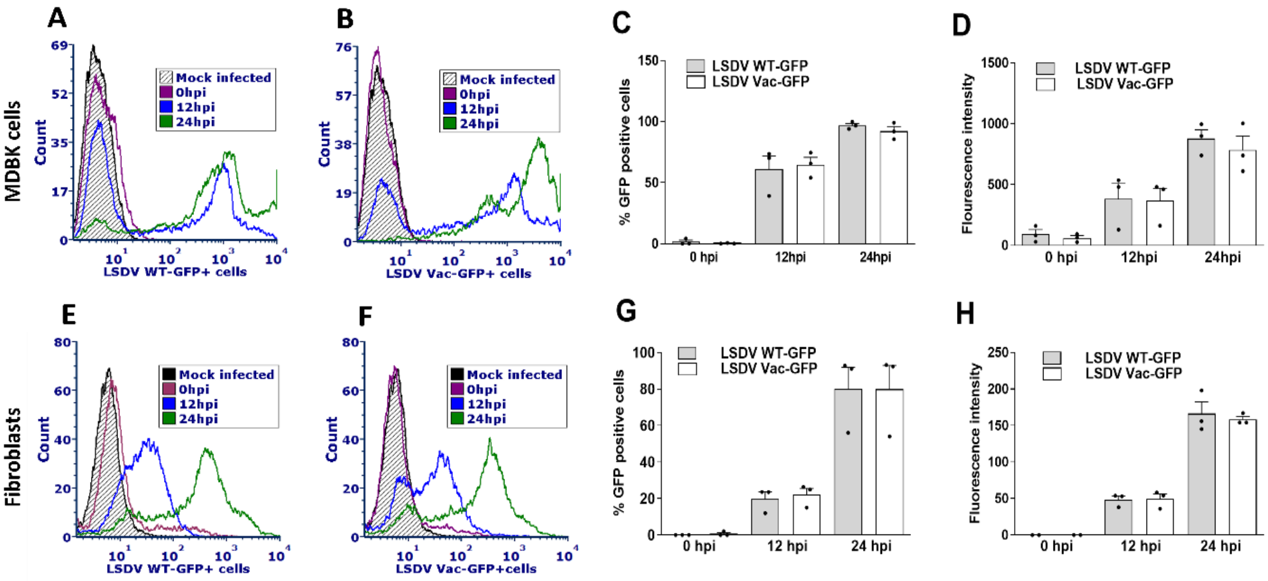


**Figure S2. GFP expression in MDBK cells and primary fibroblasts infected with GFP recombinant viruses is indicative of viral replication.**

**(A-H)** MDBK cells (A-D) and primary bovine fibroblasts (E-H) were inoculated with recombinant LSDV WT-GFP and LSDV Vac-GFP at MOI=3 for 1 h (T=0). At the time points indicated, infected cells were collected and analyzed by flow cytometry. Histograms representing LSDV WT-GFP infected MDBK cells (A, B) and fibroblast (E, F) show GFP build up in infected cells at the indicated time points. The percentage of GFP expressing cells (C, MDBK cells and G, fibroblasts cells) and the mean fluorescence intensity (D, MDBK cells and H, fibroblasts cells) are presented. Values in graphs were expressed as mean± SEM representing three biological replicates (C- D and G- H).

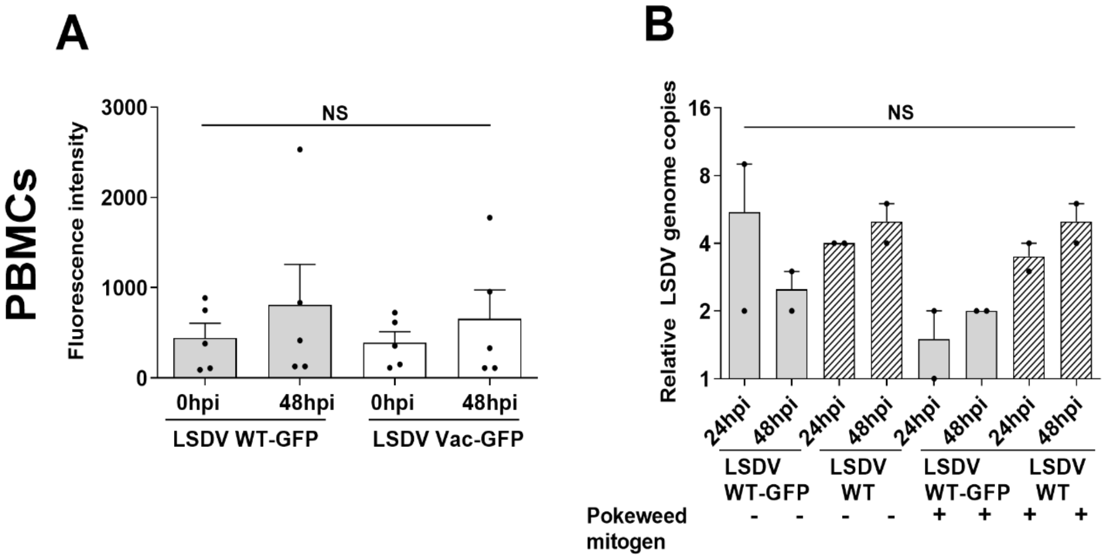


**Figure S3. Both non-stimulated and pokeweed mitogen stimulated PBMCs fail to support productive LSDV replication.**

**(A)** Non-stimulated PBMCs were inoculated with recombinant LSDV WT-GFP and LSDV VAC-GFP at MOI=3. After ~2 hours of inoculation and wash (T=0) and after 48 h cells were collected for flow cytometry analysis and the mean fluorescence intensity calculated (**this is supplementary data of Fig.2 A-C**). Bar graphs were plotted from five sets of experiments, presented as mean± SEM.

**(B)** PBMCs were either non-stimulated or stimulated with pokeweed mitogen by overnight incubation and then inoculated with LSDV strains at MOI=1 for 1h. At the times indicated, cells were collected, and viral DNA extracted and quantified by PCR. Relative genome copies were calculated as fold change to 1hpi normalized to GAPDH. Two sets were used to plot bar graph (values expressed as mean± SD).

One-way ANOVA following Tukey’s post-hoc test used to derive significance. NS- non-significant (A-B).

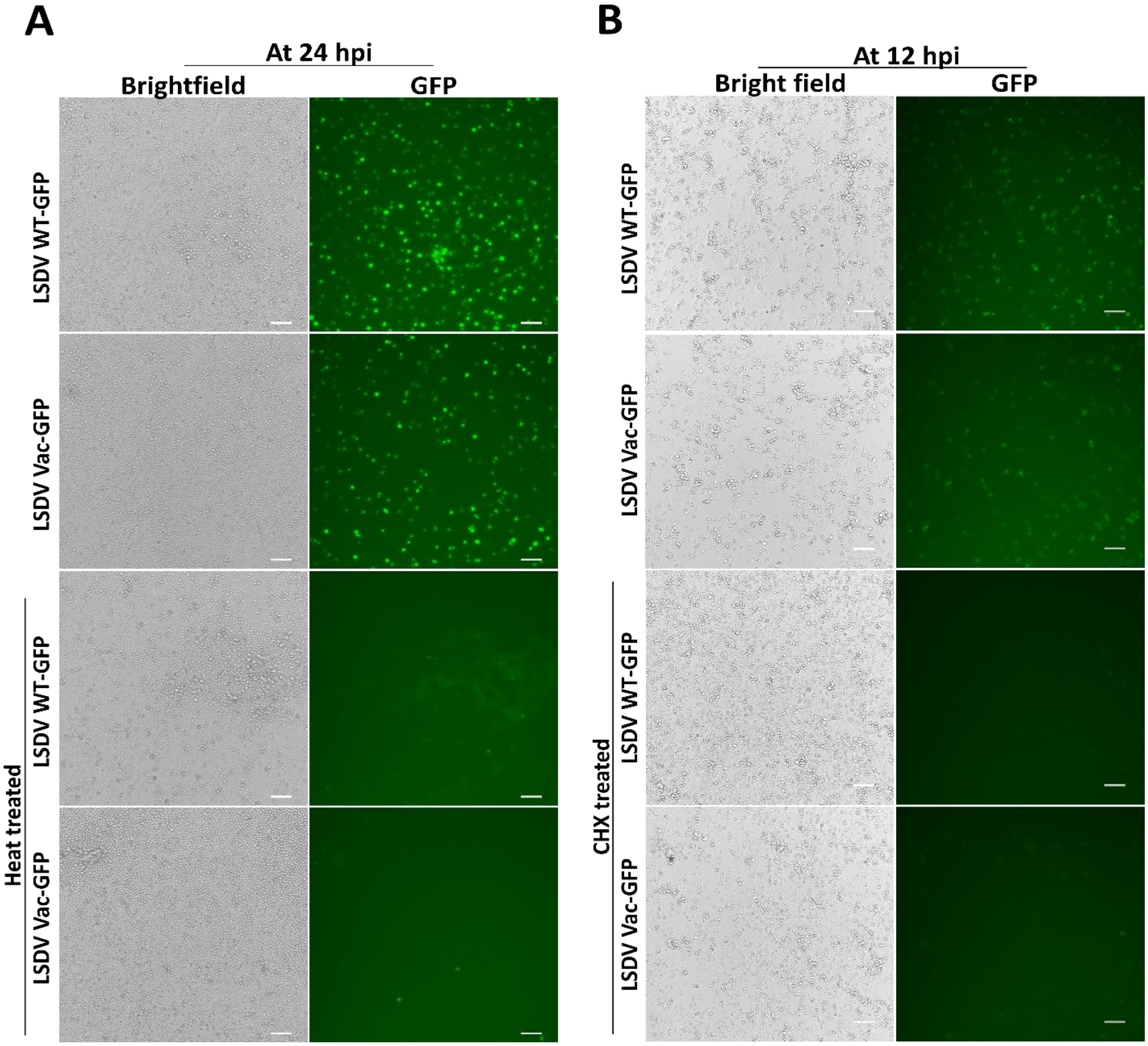
A
At 24 hpi
Brightfield
GFP
LSDV WT-GFP
LSDV Vac-GFP
Heat treated
LSDV WT-GFP
LSDV Vac-GFP
B
At 12 hpi
Bright field
GFP
LSDV WT-GFP
LSDV Vac-GFP
CHX treated
LSDV WT-GFP
LSDV Vac-GFP

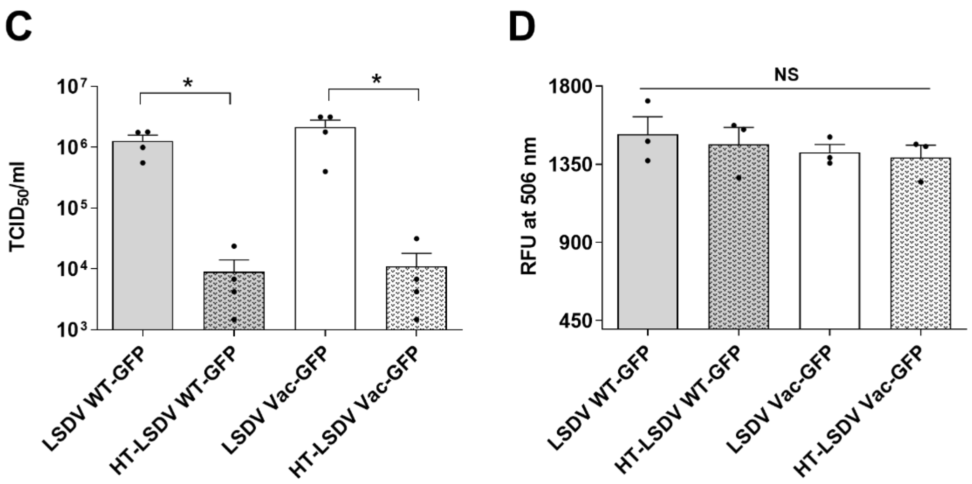


**Figure S4. *De-novo* synthesized GFP is an assay-mark of PBMCs susceptibility to LSDV infection.**

**(A)** PBMCs were inoculated with either infectious LSDV-GFP at MOI=1 or with the same volume of heat treated (HT) viruses of the same stock. After 24 hpi, cells were visualized using fluorescence microscopy. The few GFP positive cells infected with HT-GFP viruses may represent incomplete inactivation of LSDV-GFP viruses.

**(B)** PBMCs infected with LSDV-GFP viruses at MOI=1 with or without CHX pretreatment. Images taken after 12 h (B).

Representative images (A and B) from three biological repeats are presented (scale bar-100 µm)**. (C)** Viral stocks were heat incubated for 30 min at 55 ºC in a water bath. Stocks were then titrated in MDBK cells to determine any loss of infectivity. Four stocks were used to draw the figure and values presented as mean± SEM. t-test followed to compare the significance, $*p < 0.05$.

**(D)** Fluorescence intensity of preformed GFP measured in LSDV-GFP stocks (heat treated or untreated) stocks by fluorometry (excitation at 480 and emission at 506 nm) ascertaining no impact of heat treatment on GFP fluorescence. Three repeats were used to draw the histogram. Values were presented as mean± SEM. One way ANOVA was used to compare all means and post-hoc Tukey's test followed to test the significance level. NS- non-significant.

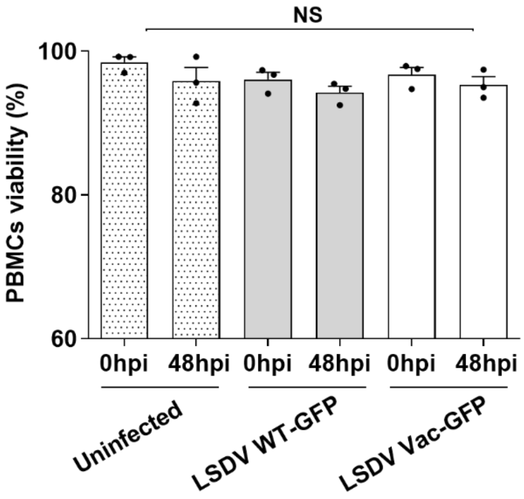


**Figure. S5. Infection with LSDV does not affect PBMCs viability.**

PBMC were either not infected or infected with GFP-LSDV strains at MOI=1 for 1 h. After 2h and 48h cells were evaluated for viability using propidium iodide (PI) exclusion assay. Values expressed as mean ±SEM representing three biological repeats. One way ANOVA was used to compare all means and post-hoc Tukey’s test followed to test significance level. NS- non-significant.

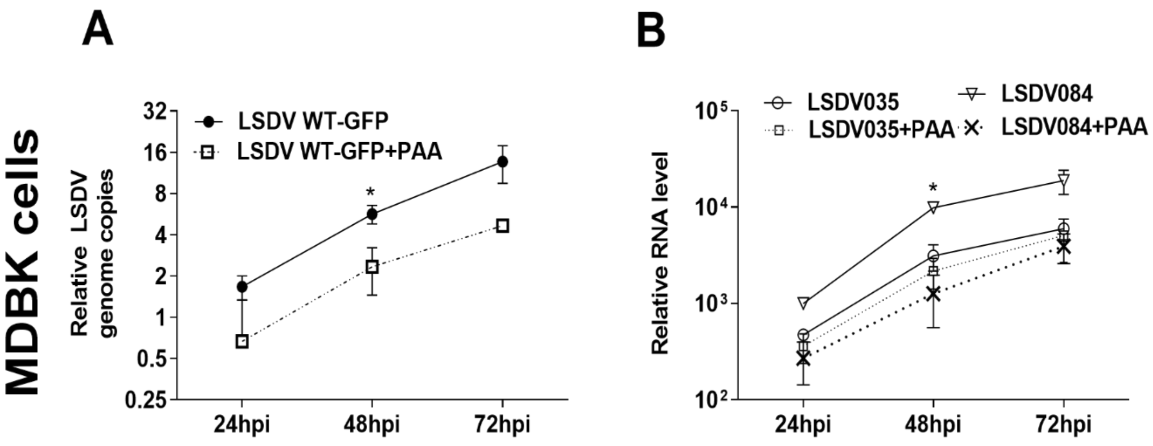


**Figure S6. LSDV084 (late gene) expression is reduced by inhibition of viral DNA replication, in contrast to LSDV035 (early gene).**

**(A-B)** MDBK cells either untreated or treated with PAA were infected with LSDV at MOI=1 for 1h and checked for inhibition of viral DNA replication (A) and viral gene expression (B) at the indicated time points. Extracted nucleic acids were used for relative quantification of viral genome copies (A) and viral RNA levels (B). Three biological sets were used to draw the graphs, presented as mean± SEM. Values represent fold change compared to 1 hpi and normalized to GAPDH DNA or RNA, respectively. Paired t-test (A) and one way ANOVA followed by post-hoc Tukey's test used to test significance. $*p < 0.05$.

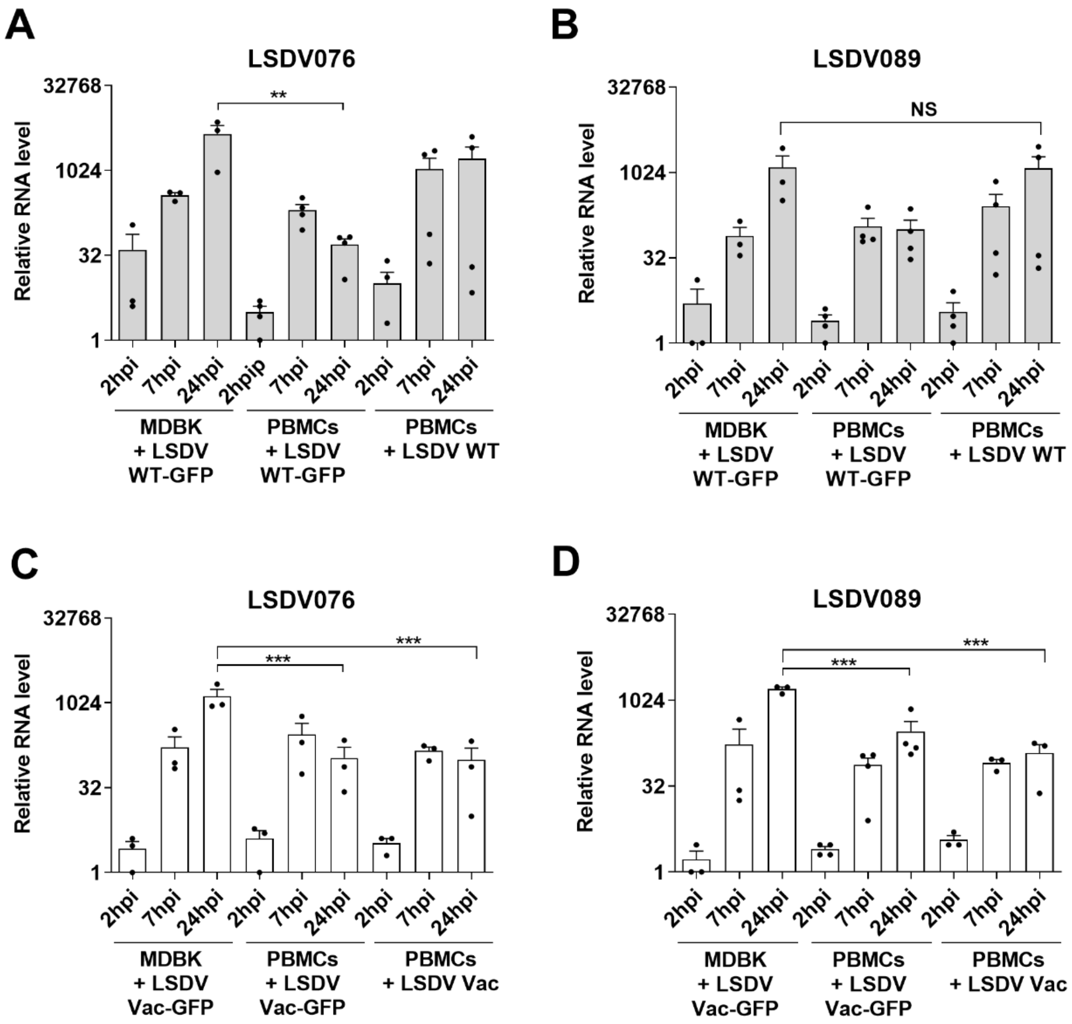
A
LSDV076
B
LSDV089
C
LSDV076
D
LSDV089
Relative RNA level
32768
1024
32
1
**
NS
***
2hpi
7hpi
24hpi
2hpip
MDBK + LSDV WT-GFP
PBMCs + LSDV WT-GFP
PBMCs + LSDV WT
MDBK + LSDV Vac-GFP
PBMCs + LSDV Vac-GFP
PBMCs + LSDV Vac

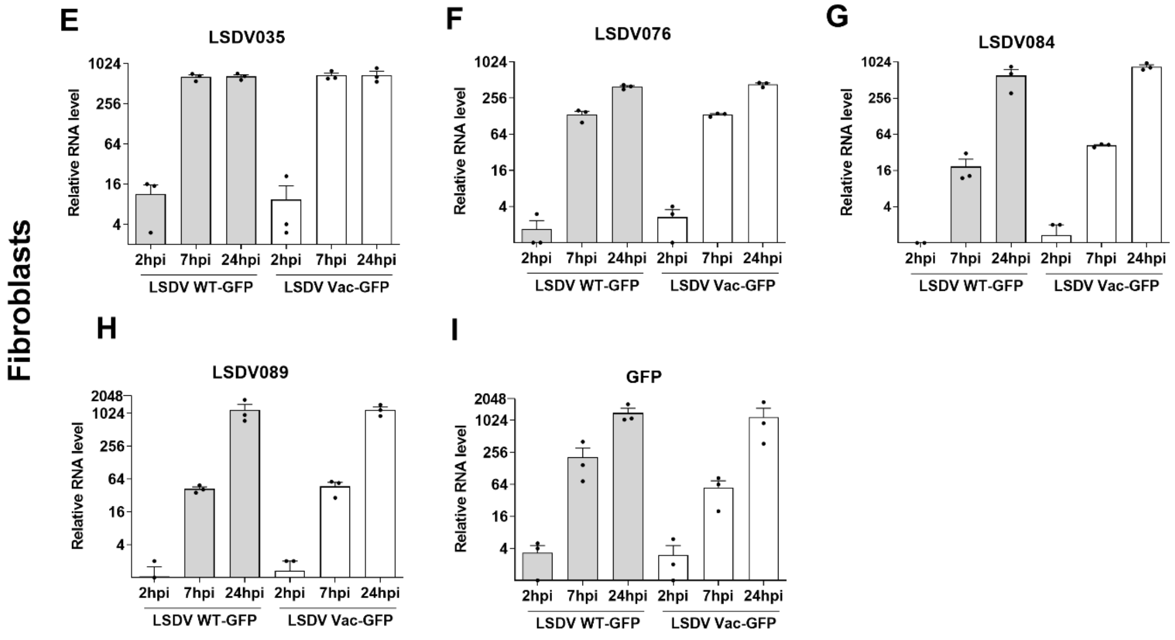


**Figure S7. LSDV genes follow a similar trend of expression in MDBK cells, PBMCs and fibroblasts.**

(**A-D**) Expression levels of LSDV076 and LSDV089 in MDBK cells and PBMCs infected with LSDV- GFP strains (Supplementary data of **Fig. 2E-J)**. Three repeats were plotted in the bar graph as mean± SEM. One way ANOVA followed by post-hoc Tukey's test used to test significance. ∗∗p < 0.01, ∗∗∗p < 0.001, NS- non-significant.

**(E-I)** Bovine foreskin fibroblasts cells were infected at MOI-1 with either LSDV-GFP strains for 1 hour. After incubation, cells were washed and replenished with fresh media. At 1h, 2h, 7h and 24h RNA was extracted. Relative changes in the levels of transcripts encoding the reporter gene GFP and viral genes were determined by RT-qPCR, normalized to GAPDH RNA (internal control) and fold change calculated against T=0 (i.e. 1 hour inoculation). Three repeats were plotted in the bar graph as mean± SEM.

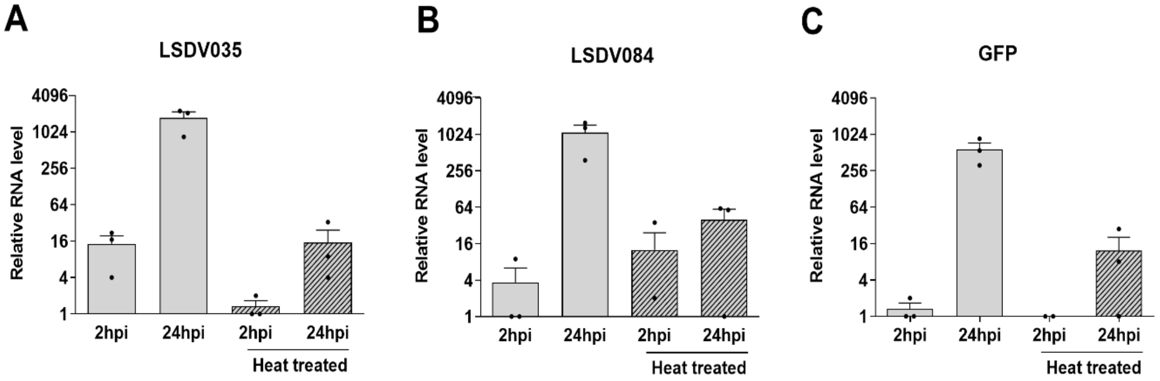


**Figure S8. Heat treatment of LSDV reduces the expression of viral genes in infected MDBK cells. (A-C)** MDBK cells were inoculated with infectious LSDV WT-GFP at MOI=1 or with the same volume of heat-treated LSDV for 1h after which cells were washed and replenished with fresh media. At 1h, 2h and 24h, cells were lysed, and RNA extracted. Relative changes in the levels of transcripts of viral genes (LSDV035 and LSDV084) and the reporter gene GFP were determined by RT-qPCR, normalized to GAPDH RNA (internal control) and fold change calculated against T=0 (i.e. 1 hour inoculation). Values were presented as mean± SEM.

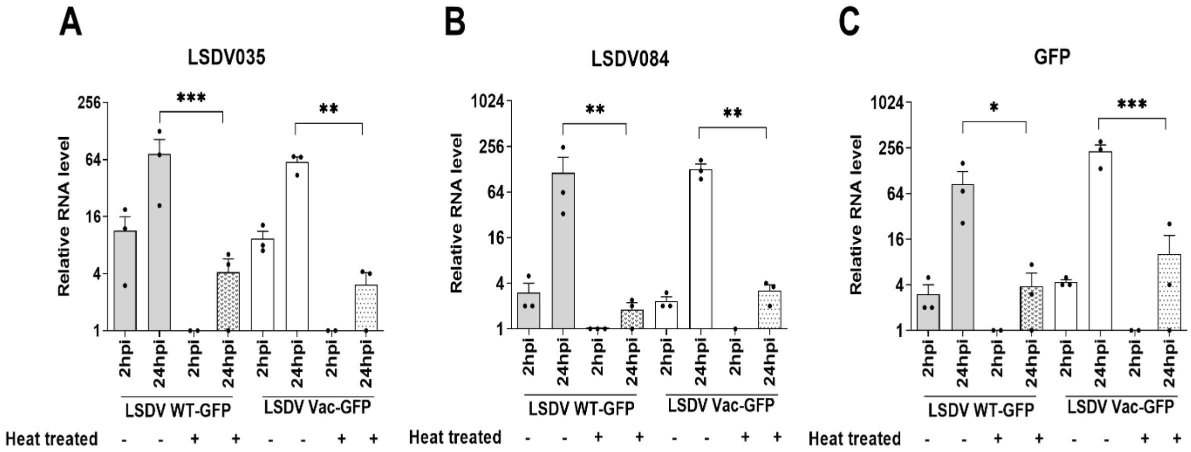


**Figure S9. Heat treatment of LSDV reduces the expression of viral genes in inoculated PBMCs. (A-C)** PBMCs were inoculated with infectious LSDV WT-GFP at MOI=1 or with the same volume of heat-treated LSDV for 1h after which cells were washed and replenished with fresh media. At 1h, 2h and 24h, cells were lysed, and RNA extracted. Relative changes in the levels of transcripts of viral genes (LSDV035 and LSDV084) and the reporter gene GFP were determined by RT-qPCR, normalized to GAPDH RNA (internal control) and fold change calculated against T=0 (i.e. 1 hour inoculation). Values were presented as mean± SEM. One -way Anova with Sidak multi-comparison test used to measure significance level. ∗p <0.05, ∗∗p <0.01 and ∗∗∗p <0.001.

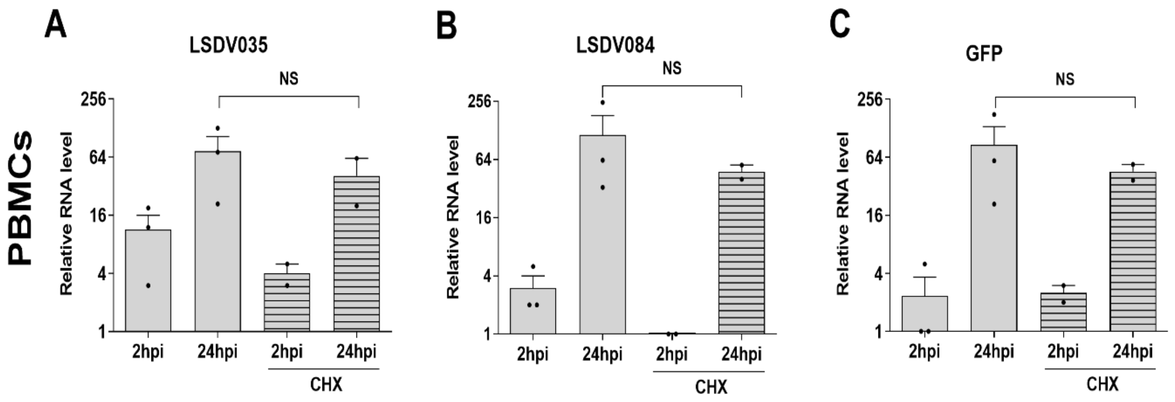


**Figure S10. The levels of LSDV transcripts are only mildly affected by CHX treatment in infected PBMCs.**

**(A-C)** PBMCs either untreated or treated with CHX were infected with LSDV WT-GFP at MOI=1 for 1h after which cells were washed and replenished with fresh media. At 1h, 2h and 24h, cells were lysed, and RNA extracted. Relative changes in the levels of transcripts of viral genes (LSDV035 and LSDV084) and the reporter gene GFP were determined by RT-qPCR, normalized to GAPDH RNA (internal control) and fold change calculated against T=0 (i.e. 1 hour inoculation). Values are shown as mean± SEM of three biological repeats. One way ANOVA followed by post-hoc Tukey's test used to test significance. NS- non-significant.

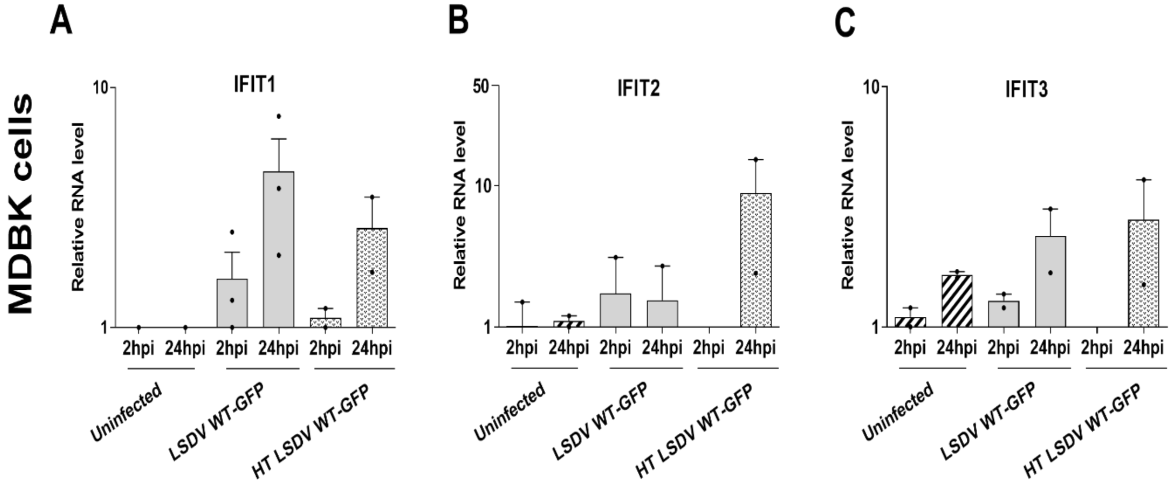


**Figure S11. MDBK cells infected with LSDV show insignificant changes in ISGs' RNA levels.** **(A-C)** MDBK cells were infected with LSDV-GFP viruses at MOI=1 for 1h after which cells were washed and replenished with fresh media. At 1h, 2h and 24h, cells were lysed, and RNA extracted. Relative changes in the levels of IFI1-3 transcripts were determined by RT-qPCR, normalized to GAPDH RNA (internal control) and fold change calculated against T=0 of uninfected cells. RNA used here were from the experiment described in **Fig. 2E-J**. Values are shown as mean± SEM of three biological repeats.

### Table S1

| Sl. no | Primer sets | Sequence (5'-3') | Amplicon (bp) | Notes/Remarks/Gene ID |
|---|---|---|---|---|
| 1. | LSDV035F | CCATCTCGCGCTAGATGAATAC | 149 | 27% homology to VV gene (E5R) under an early promoter |
| | LSDV035R | TGTATCCTAGCTTTTTCGGAGG | | |
| 2. | LSDV076F | AGGTAGCGGACGAAATCTTTCC | 93 | 37 % homology to VV gene-H5R under an early promoter |
| | LSDV076R | CCGCTTTCTTACGAACAACTGG | | |
| 3. | LSDV084F | AATGGGTCACAAGGGACAAATC | 137 | 81% homology to VV gene-D6R under a late promoter |
| | LSDV084R | TGGCTCCCATCATCATTTGTTG | | |
| 4. | LSDV089F | TTGGACATTGTTGGCATCCTTC | 115 | 74% homology to VV gene-D12L under E/L promoter |
| | LSDV089R | AGATAGTGCGATTATGGTAGCG | | |
| 5. | GFP-F | CACCATCTTCTTCAAGGACGAC | 212 | Expressed under vaccinia synthetic early/late promoter |
| | GFP-R | TCTTGAAGTTCACCTTGATGCC | | |
| 6. | GAPDH-F(RNA) | AGGTCGGAGTGAACGGATTC | 197 | Self-design, NM_001034034.2 |
| | GAPDH-R(RNA) | CATTGATGACGAGCTTCCCG | | |
| 7. | GAPDH-F(DNA) | GTGATGCTGGTGCTGAGTAT | 139 | NC_037332, (Wang et al., 2017) |
| | GAPDH-R(DNA) | GCTCTCACATTCCTAAGTCC | | |
| 8. | CAPRI-F | AAATGAAACCAATGGATGGGATA | 89 | (Bowden et al., 2008) |
| | CAPRI-R | AAAACGGTATATGGAATAGAGTT | | |
| 9. | LSDV05-F | AAGGCTATGGGAGAGTTTG | 973 (Pair-A) | Self-design, LSDV005 and LSDV006 |
| | LSDV06-R | CGATGATAGGATACAGAG | | |
| 10. | EGFP-F | AAGGCTATGGGAGAGTTTG | 243 (Pair-B) | Self-design, pEGFP-N1 and LSDV006 |
| | LSDV06-R | GATGAACTTCAGGGTCAGC | | |
| 11. | LSDV05-F | AACGAGAAGCGCGATCAC | 238 (Pair-C) | Self-design, LSDV005 And pEGFP-N1 |
| | EGFP-R | CGATGATAGGATACAGAG | | |
| 12. | LSDV05-F1 | **GCTCCCGGCCGCCATGG*CCGCGG*** GATATGAAAACAAACACAAAAATAATAC | | Self-design, LSDV005, (Letters in italics show restriction digestion sites, bold letters represent pGEMT terminal sequence and bold underlined letters showing vaccinia derived promoter sequence) |
| | LSDV05-R1 | **<u>TATTTATATTCCAAAAAAAAAAAATAAAATTTCAATTTT</u>***GAATTC*TTACTTTTTAACTATGTACTTTTC | | |

| | | | | |
|---|---|---|---|---|
| 13. | EGFP-F | AGTACATAGTTAAAAAGTAAAAAA TTGAAATTTTATTTTTTTTTTTGGA ATATAAATAATGGTGAGC AAGGGCGAG | | Self-design, pEGFP-N1 |
| | EGFP-R | TTGTAGTTTTATAA*CTCGAG*TTACT TGTACAGCTCGTCCATGC | | |
| 14. | LSDV06-F2 | *CTCGAG*TTATAAAACTACAACATTA TAG | | Self-design, LSDV006 (Letters in italics show restriction digestion site and bold letters represent plasmid terminal sequences) |
| | LSDV06-R2 | **CAGGCGGCCGCACTAGTG***ACTAG T*AAATGAAAGTTATAAATTTTATC | | |
| 15. | IFIT1-F | GGAACGTGCTGTGCAACTAA | 136 | XM_010819765.1, (Cheng et al., 2017) |
| | IFIT1-R | TTTGTCGAGTGCTTTCATGC | | |
| 16. | IFIT2-F | ACCCCATTAACCCTTTGAGG | 249 | BT025389.1, (Stepke et al., 2018) |
| | IFIT2-R | TGTTGGGCATGCATTTTAGA | | |
| 17. | IFIT3-F | TGCTGACAAGGTGAAACGAG | 111 | NM_001075414, (Cheng et al., 2017) |
| | IFIT3-R | TTTTTCCCACCGCACTTTAC | | |
| 18. | IFITM3-F | GTGGCATTCGCCTACTCTGT | 142 | NM_001078141.2, (Stepke et al., 2018) |
| | IFITM3-R | CGATGAGGACGACAGTCAGA | | |
| 20. | ISG15-F | AGAAGATCAATGTGCCTGCTTT | 161 | NM_174366.1, (Cheng et al., 2017) |
| | ISG15-R | CTTGTCGTTCCTCACCAGGAT | | |
